# Forecast Padding Enhances Accuracy and Robustness of EEG-Phase-Synchronized TMS

**DOI:** 10.1101/2025.03.19.643976

**Authors:** Yu-Cheng Chang, Pin-Hsuan Chao, Yan-Ming Kuan, Chiu-Jung Huang, Li-Fen Chen, Wei-Chung Mao, Tung-Ping Su, Sin-Horng Chen, Chun-Shu Wei

**Author notes:** The corresponding authors are C.-S. Wei and W.-C. Mao.

## Abstract

Closed-loop neuromodulation is a promising personalized treatment for various neuropsychiatric disorders, delivering precise stimuli based on real-time brain signals. However, its clinical potential is currently limited by technical challenges inherent in its real-time nature. This article addresses the primary technical challenges in EEG-Phase-Synchronized Transcranial Magnetic Stimulation (TMS), including poor stimulation accuracy and inefficient biomarker detection (correlated with deadlock). These challenges arise from the vulnerability of existing algorithms to filter edge effects. Inspired by predictive coding theory in neuroscience, we propose a novel signal padding method (forecast padding) to mitigate the filter edge effect. To properly quantify the improvements that forecast padding brings about in real-world systems, we introduce a novel delay-relevant validation framework and demonstrate its reliability using experimental data from a real system. Through this framework, we demonstrate that forecast padding significantly improves both stimulation accuracy and deadlock rate. Given the pervasive impact of filter edge effects in closed-loop neuromodulation and other signal processing domains, forecast padding shows broad application potential across various fields.

## I. Introduction

Closed-loop neuromodulation, which dynamically adjusts stimulation parameters based on real-time physiological feed-back, has emerged as a promising approach in treating neuropsychiatric disorders. By synchronizing stimuli with predefined neural biomarkers (targets), this method can effectively improve therapeutic outcomes and reduce both inter- and intrasubject variability [1], [2], [3]. However, technical challenges

The authors gratefully acknowledge Bo-Shan Wang, Pei-Chun Chang, Yong-Sheng Chen, Chi-Yen Hsieh, Yu-Feng Liu, Hsin-Yu Feng, Joey Chen, Lisa Chiu, Max Chiang, and Raymond Chou for their assistance in providing access to TMS and EEG equipment and offering valuable technical support. This work was supported in part by the National Science and Technology Council (112-2222-E-A49-008-MY2, 112-2321-B-A49-012, 113-2321-B-A49-019, and 113-2124-M-001-020), the Higher Education Sprout Project of National Yang Ming Chiao Tung University (NYCU) and the Ministry of Education of Taiwan, and Cheng Hsin General Hospital (CY11103, CY11205).

Y.-C. Chang and S.-H. Chen were with the Dept. of Electronics and Electrical Engineering, Y.-M. Kuan was with the Dept. of Computer Science (CS), P.-H. Chao is with the Inst. of Biomedical Engineering (BME), L.-F. Chen and C.-J. Huang are with the Inst. of Brain Science, and C.-S. Wei is with the Dept. of CS and the Inst. of BME, NYCU, Hsinchu, Taiwan; W.-C. Mao, and T.-P. Su are with the Dept. of Psychiatry, Cheng Hsin General Hospital, Taipei, Taiwan. The corresponding authors are C.-S. Wei and W.-C. Mao. continue to hinder its broader clinical application. This article primarily addresses the challenges of a type of closed-loop neuromodulation, EEG-phase-synchronized transcranial magnetic stimulation (TMS) [4], [5], [6], which delivers magnetic stimuli based on the phase of EEG rhythm. Our focus on this system is twofold: first, due to its therapeutic potential for depression [7], [8], [9], [10], and second, because its demanding temporal precision requirement (typically within milliseconds) exemplifies the common technical challenges encountered in closed-loop neuromodulation more broadly.

### A. Primary Technical Challenges

To systematically address these technical challenges, we categorize them into three main aspects (Readers unfamiliar with the technical aspects of closed-loop TMS are encouraged to consult the Preliminary Sections in the Supplementary Information):

1. **Predictive Performance**: Since the EEG phase must be extracted in real-time, phase prediction can only use EEG data up to the current time. This fundamental limitation leads existing algorithms [11], [12], [13] to exhibit significant deviations from the ground truth (acausally estimated benchmark, defined in Preliminary Section 2).
2. **Robustness**: Several studies have reported unusually long target identification times (communication deadlock) in their systems [14], [7], [15], with limited prior understanding of this phenomenon. Our analysis reveals that this results from unbalanced phase prediction (UBPP) induced by the filter edge effect (see Sections I-B.1 and III-C.2). Simply put, the highly concentrated predicted phases prevents the target phase from being reached. The deadlock issue greatly limits the clinical utility of the system, hindering applications that require a sufficient pulse rate, such as phase-synchronized rTMS/TBS protocols [10], [7].
3. **Replicability**: Incomplete consideration of system delays has led to two critical issues in the field: overrated stimulation accuracy and irreproducible phase-dependent modulatory effects [5], [4]. A detailed comparison of these issues across different systems is provided in Table 1 of the Preliminary Section 3.4.

### B. Causality Limitation

Although the above three challenges may seem superficially independent, they share a common root cause: the causality limitation (the inherent inability to access future information) [12] (see Figure 1). This limitation fundamentally dictates that, due to the nature of online scenarios or the obscuring effect of the TMS artifact in offline settings, the phase at any given time can only be reliably estimated from past and present signals; the ground truth, therefore, remains inaccessible.

**Fig. 1:**
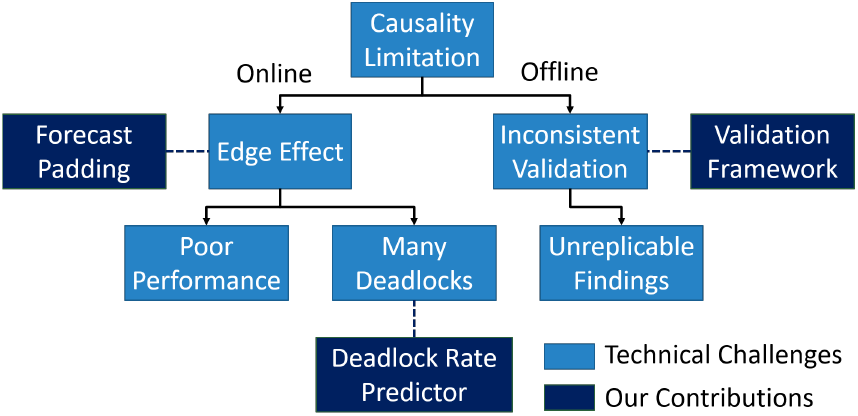
Causality linitation is the root cause of primary technical challenges.

#### 1) Causality Limitation in Online Stimulation

During realtime stimulation, zero-phase filtering (a bidirectional filtering technique requiring both past and future data) is employed in most existing phase prediction algorithms [16], [12], [11], [14] to extract the rhythm of interest. This process involves padding zeros in place of unseen future data to maintain output dimensions. However, convolving the kernel with these padded zeros causes distortion and information loss near the splice joint (t=0), as illustrated in Figure P4a of Preliminary Section 4. This distortion leads to biased predictions towards 90° and 270° (see Preliminary Section 4). Beyond reducing accuracy, the concentrated predictions increase the risk of missed detections and, if persistent, can result in communication deadlocks (see Section III-C.2). To address these limitations, this study introduces a novel padding technique, forecast padding, to overcome the problems caused by conventional zero padding.

#### 2) Causality Limitation in Offline Validation

In our review of existing validation methods for quantifying stimulation accuracy, we found diverse and incomplete considerations of delay across studies, leading to unconvincing accuracy reports (summarized in Table 1 of Preliminary Section 3.4). The underlying cause is that the TMS artifact obscures the signal around the stimulation onset, preventing the calculation of ground truth at the pulse location and complicating accurate quantification. This limitation has led researchers to employ alternative validation methods to circumvent the pulse occurrence (in-silico and fake-pulse methods; see Preliminary Section 3). However, these methods often overlook critical system delay properties, resulting in overrated stimulation accuracy. Specifically, most methods either ignore or underestimate the transport delay (*τ*_*all*_) of the closed-loop system. Some studies report algorithms’ in-silico predictive performance [11], [13], [17] but these results fail to reflect the actual stimulation accuracy of a closed-loop system, which is influenced by both the algorithm’s predictive performance and delay jitter in online scenarios. To address these limitations, we integrated existing methods with our previously proposed delay model [18] into a delay-relevant validation framework. This framework notably enables the use of in-silico simulation to objectively evaluate the stimulation accuracy of actual closed-loop systems, providing a practical method for system development and validation.

### C. Main Contributions

In conclusion, to address the challenges imposed by causality limitations, this paper presents three main contributions:

1. Establishing the relationship among UBPP, filter edge effect, and system deadlock.
2. Developing a framework to estimate the accuracy and robustness of closed-loop TMS systems.
3. Introducing a forecast padding technique that can be applied to various algorithms to improve system accuracy and reduce deadlock occurrence.

## II. Material and Methods

The section is organized as follows: first, we present the forecast padding technique and its theoretical foundation; second, we present both online and offline experiments to demonstrate the improvements in accuracy and robustness achieved through forecast padding; and finally, we investigate the relationship among filter edge effect, UBPP, and system deadlock. The EEG data used in each experiment are documented in Supplementary Information S.4.

### A. Predictive Coding and Forecast Padding

As explained previously, zero padding discards future information by assuming zero input for unseen data, which contradicts the information processing mechanisms of the human brain. The brain’s ability to predict incoming data before actual reception is well-established in neuroscience through predictive coding theory [19], [20], [21], [22]. This theory, originating from Helmholtz’s concept of unconscious inference in 1867 [23], posits that the brain actively predicts sensory input based on prior experiences rather than passively receiving information. The theory reached a significant milestone with Rao’s hierarchical model in the late 1990s [24], which interprets perception as an interplay between top-down prediction and bottom-up sensory processes. According to the model, during the top-down process, the higher level of the cortex predicts lower level’s sensory input; in the subsequent bottom-up process, the prediction is then compared with the actual sensory input to update the predictive model at the higher level. Spratling later demonstrated that this model is mathematically equivalent to the biased competition model, where top-down processing acts as a selective attention mechanism to accentuate the relevant sensory information [25]. In essence, predictive coding operates on the principle that “what we predict directs what we sense, and what we sense updates what we predict.”

Drawing upon this biological principle of predictive processing, we introduce a novel padding scheme called forecast padding to address the filter edge effect. The correspondence between predictive coding and forecast padding is shown in Figure 2a, where the kernel can be analogously linked to the sensory encoder at the lower level, and the predictive algorithm to the predictive estimator at the higher level. As illustrated in Figure 2b, the overall computation process involves two convolutions: the first being the bottom-up process and the second the top-down process.

**Fig. 2:**
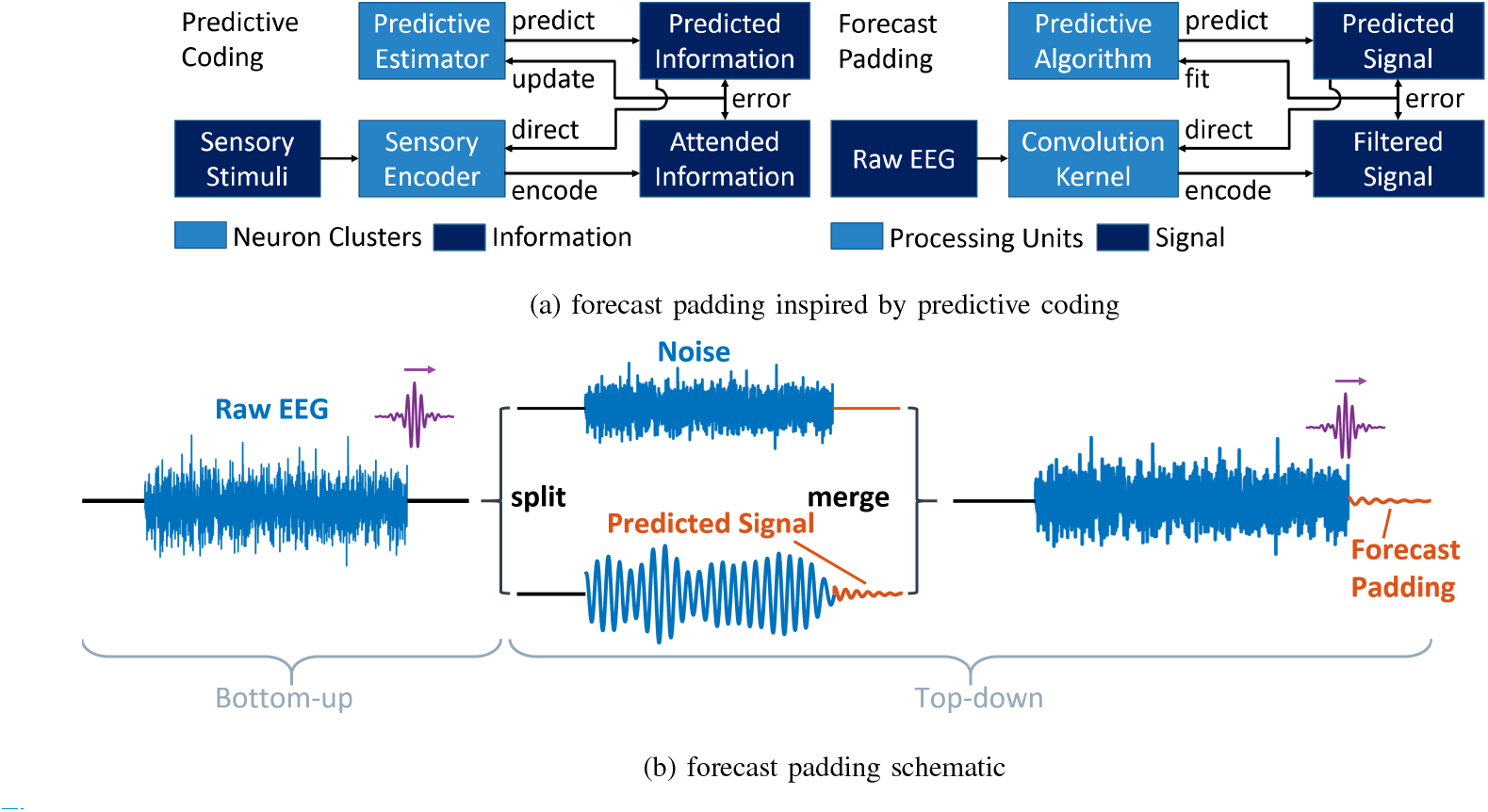
(a) Correspondence between the elements of predictive coding and forecast padding. (b) Forecast padding preserves task-related information near t=0 by augmenting the raw signal with a synthesized segment, which combines a noise baseline and the t*>*0 portion of the predicted signal. This approach maintains signal continuity while preserving critical information at the temporal boundary.

#### 1) Implementation Process

In the first convolution, the initially zero-padded raw signal is convolved with a kernel to separate it into signal (task-related part) and noise (non-task-related part), which are then padded separately. The signal is padded with its predicted values, representing task-related information. Conversely, the noise is padded with its baseline level (DC offset) to suppress its power. Unlike zero-padding, which causes a sharp transition and ringing after filtering, the noise padding ensures smooth concatenation and eliminates noise power simultaneously. Finally, the sum of signal and noise paddings is concatenated back to the raw data to replace the padded zeros. The second convolution is then applied to the concatenated data. As the kernel slides across the splice joint (t=0), it now combines information from both the raw EEG and the predicted signal, allowing the convolution to more effectively emphasize task-related information and suppress irrelevant noise.

#### 2) Application to EEG Phase Prediction

Forecast padding is self-adaptive to various downstream tasks, as the kernel can be dynamically adjusted to meet the specific task requirements. When applied to EEG phase prediction, as in this paper, the kernel functions as a bandpass filter to extract the sensorimotor *µ* rhythm. The *µ* rhythm was chosen for two reasons: first, its relatively stable oscillatory characteristics provide an ideal testbed for evaluating the phase prediction algorithm [12]; second, it serves as a crucial neurophysiological marker in understanding phase-dependent corticospinal excitability and plasticity [4]. An algorithm is then employed to predict its future oscillation. The predicted signal is shifted to the raw EEG’s baseline and appended to it. Since the bandpass signal does not contain a DC component and the baseline level of the raw data is attributed solely to noise, the baseline shifting is equivalent to the noise padding described previously. Finally, the concatenated data is filtered again to generate a less distorted *µ* rhythm, leading to a more accurate phase prediction. The implementation details of forecast padding in each algorithm are described in Supplementary Information S.10.

### B. Predictive Performance and Robustness Evaluation

The mean absolute error (MAE) and the imbalance factor (IBF) are used to quantify predictive performance and robustness of the algorithms, respectively. The MAE is defined as follows:

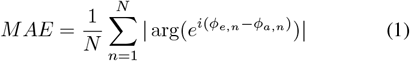

Here, *N* is the number of EEG epochs, *ϕ*_*e,n*_ is the phase estimated by the algorithms, and *ϕ*_*a,n*_ is the ground truth. To quantify the UBPP phenomenon, which is the primary cause of deadlock, we developed the imbalance factor (IBF). This metric allocates the predicted phases to *N*_*b*_ bins in a polar histogram, each covering a sector of 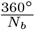. The IBF is defined as follows:

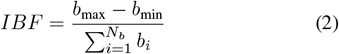

Here, *b*_*i*_ represents the bin count of the *i*^th^ bin, *b*_max_ is the maximum bin count, and *b*_min_ is the minimum bin count. IBF ranges from 0 to 1, where 0 means that phases are uniformly distributed across all sectors, while 1 means that all phases belong to the same sector. In this study, *N*_*b*_ = 20, providing sufficient angular resolution (18° per sector) to capture the phase distribution patterns while ensuring adequate sample sizes per bin for computing IBF.

Using the EEG recorded from the fake-pulse experiment, we evaluated eight phase prediction algorithms offline. For all algorithms, we used MATLAB’s ‘filtfilt’ function for filtering, which implements Gustafsson’s method [26]. This approach applies zero padding while optimizing initial conditions to minimize filter edge effects, representing a standard practice in signal processing. The algorithms evaluated include:

1. Four base algorithms (non-FP condition: using ‘filtfilt’’s built-in initial condition optimization):
  - AR (Autoregressive) [11]
  - ETP (Educated Temporal Prediction) [12]
  - FFT (Fast Fourier Transform) [17]
  - PR (A newly proposed algorithm representative of conventional linear phase prediction approaches)
2. Their forecast padding counterparts (FP condition: applying forecast padding before filtering with ‘filtfilt’):
  - ARFP (AR with Forecast Padding)
  - ETPFP (ETP with Forecast Padding)
  - FFTFP (FFT with Forecast Padding)
  - PRFP (PR with Forecast Padding)

We assessed their predictive performance using MAE and robustness using IBF. Both MAE and IBF were calculated using the phases computed at t=0 for all algorithms to ensure consistency in evaluation. To assess the statistical significance of improvements after applying forecast padding, we used a right-tailed Wilcoxon signed-rank test, which is suitable for paired comparisons of non-normally distributed data.

### C. Online Experiments

We evaluated the efficacy of forecast padding in a realworld online closed-loop TMS system by assessing stimulation accuracy and deadlock rate. Each participant (N = 20) underwent five phase-synchronized stimulation sessions: three fake-pulse sessions testing five algorithms (AR, ETP, FFT, PR, and ARFP) and two real-pulse sessions comparing AR and ARFP. AR was selected for forecast padding integration due to its computational efficiency and widespread adoption in realtime phase prediction applications. This study was approved by the Institutional Review Board of Cheng Hsin General Hospital on January 4th, 2023 (CHGH-IRB No. (985)111A-63). Detailed experimental procedures and data preprocessing pipelines are presented in Supplementary Information S.11 and S.12, respectively.

### D. Validation Framework for Stimulation Accuracy

A novel validation framework is proposed to comprehensively quantify stimulation accuracy in online scenarios through three commonly used methods: in-silico (computational simulation), fake-pulse (hardware simulation without actual stimulation), and real-pulse validation (actual TMS delivery). All three methods are modified to consider both the mean and inter-trial variance (jitter) of delay. To demonstrate the reliability of the framework, statistical tests (detailed in Supplementary Information S.6) were conducted to verify the equivalence of accuracies reported by all three methods. For in-silico and fake-pulse validation, MAE is chosen as the evaluation metric for accuracy because it simultaneously accounts for both bias (the distance between the mean pulse phase and the target phase) and variance (the spread of the pulse phase). For real-pulse validation, a qualitative approach is proposed to visualize the pulse phase.

#### 1) In-Silico Validation

We fitted the distribution of *τ*_*sys*_ using delay measurements from the real-pulse session [27]. *τ*_*algo*_ was treated as constant due to its significantly lower variation compared to *τ*_*sys*_ (see Supplementary Information S.1 and S.3). Consequently, the simulated pulse location *τ*_*all*_ for each trial can be approximated as the sum of a constant 𝔼 (*τ*_*algo*_) and a randomly sampled *τ*_*sys*_. The best-fitting distribution for *τ*_*sys*_ was determined to be a Weibull maximum distribution with parameters (shape=279.45, location=1824.96, scale=1796.68). The EEG data and the trigger information of the fake-pulse sessions were used in the in-silico validation. For each simulated pulse, the 2-second C3-laplacian signal preceding t=0 was used as the input to the five algorithms. To evaluate the performance of the algorithms at different levels of delay, we adjusted the location and scale parameters to achieve different mean *µ* and variance *σ*^2^ of *τ*_*all*_. As illustrated in Figure 4h, the simulated pulse locations *{τ*_*all,m*_*}* _*m*=1,2,…,*M*_ were randomly sampled from the Weibull maximum distribution (c=279.45, mean=*µ*, SD=*σ*), where *M* = 1000 in this study. During the sampling procedure, the negative part of the distribution was truncated to ensure physical validity.The MAE for trial *n* is defined as

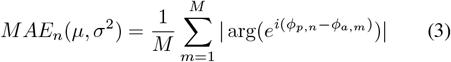

where *ϕ*_*p,n*_ is the predicted phase for trial *n* at *t* = *µ* and *ϕ*_*a,m*_ is the ground truth at *t* = *τ*_*all,m*_. The MAE averaged across trials is

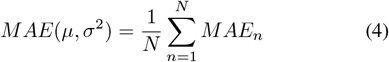

#### 2) Fake-Pulse Validation

In fake-pulse validation, we analyzed the EEG data and trigger information recorded in the fake-pulse sessions where the TMS output is replaced by the event markers. The best-fitting distribution of 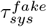 was found to be a power-log-normal distribution (c=1.83, s=0.21, location=-9.47, scale=39.03) [27]. To account for the different distributions between *τ*_*all*_ and 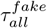 while maintaining focus on actual TMS accuracy, we developed a distribution adjustment method based on the following theorem:

##### Theorem 2.1

Let *H* : ℝ *→* [0, 1] be the cumulative distribution function (CDF) of the source distribution (powerlog-normal in this paper), and *G* : [0, 1] *→ ℝ* be the quantile function (inverse CDF) of the target distribution (Weibull maximum in this paper). Let denote *∘* function composition. Then the adjusted marker location

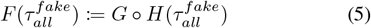

follows the target distribution.

*Proof* By the *probability integral transform*, 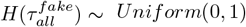 *Uniform*(0, 1). Let *T*_*X*_ (*x*) be the CDF of the target distribution. The *inverse transform sampling* states that for any *U ~ Uniform*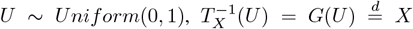. Therefore, 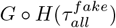 follows the target distribution.

This transformation preserves the monotonic relationship between 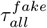 and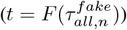, which is essential given that the fake-pulse system shares most components with the real-pulse system except for the stimulator. The performance of the algorithm for the trial *n* is given by

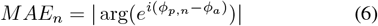

where *ϕ*_*p,n*_ is the real-time predicted phase at *t* = 𝔼 (*τ*_*all*_) for trial *n* and *ϕ*_*a*_ is the offline-calculated ground truth at the adjusted marker location 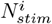.The MAE averaged across trials is

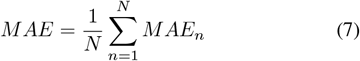

. To visualize the phase locking performance of each algorithm, polar histogram was used to illustrate in the distribution of the ground truth phase at the adjusted marker locations.

#### 3) Real-Pulse Validation

Real-pulse validation provides a graphical evaluation of stimulation accuracy by visualizing the aggregate pre-pulse waveform. The process is similar to calculating the event-related potential, except that the signal is filtered before averaging due to the significant 60-Hz line noise in our data. The validation process consists of the following six steps:

1. Epoch Extraction: The EEG data were divided into 4-second epochs, spanning from *t* = *t*_*s*_ *−* 1999 ms to *t* = *t*_*s*_ + 2000 ms, where *t*_*s*_ denotes the pulse onset.
2. Filtering: A linear-phase causal FIR bandpass filter (passband: 8–13 Hz, order: 140, group delay (*grd*): 70) was applied to each epoch. This shifts the pulse to *t* = *t*_*s*_ + *grd*.
3. Standardization: The filtered epochs were standardized by subtracting the mean and dividing by the standard deviation of the interval [*t*_*s*_ *−*100 ms, *t*_*s*_ *−*1 ms].
4. Dominant Frequency Determination: A 1024-point FFT was computed for the interval [*t*_*s*_ *−*1024 ms, *t*_*s*_ *−*1 ms] to determine the dominant frequency (the frequency bin with the largest magnitude response in the alpha band) for each epoch. The *µ* rhythm cycle, *T*_*µ*_, is the reciprocal of this dominant frequency.
5. Time Axis Normalization: To unify the time axis of each epoch for subsequent averaging, the origin was recentered at the new pulse location (*t*_*s*_ + *grd*) and the axis unit was normalized to “the number of *T*_*µ*_”.
6. Phase-Locking Evaluation: The average waveform at *t* = *−* 1 (unit: *T*_*µ*_) was inspected to examine the phaselocking performance. We examine one cycle preceding the pulse to avoid the ringing artifact around the TMS pulse induced by filtering.

### E. Validation Framework for Robustness

#### 1) Filter Edge Effect and UBPP

To explore the relationship between filter edge effect and UBPP, we scrutinize the computational process of AR and its forecast-padded counterpart, ARFP, as an example. The computational process consists of three intermediate steps:

1. Apply zero-phase bandpass filtering to the EEG epochs.
2. Remove the distorted edge portion and fit the autoregressive model to forecast the future waveform. For ARFP, append the forecast waveform to the raw EEG and apply bandpass filtering again.
3. Apply the Hilbert transform to the resulting narrow-band signal to calculate the instantaneous phase at t=0.

We illustrate the waveforms from steps 1 and 2, along with the benchmark signal obtained by zero-phase filtering the continuous EEG and extracting the corresponding time intervals of the EEG epochs. Due to computational constraints, we randomly selected 100 prediction attempts per algorithm from each of 10 randomly chosen fake-pulse sessions, resulting in a total of 1000 EEG epochs for visualization. Through this visualization, we demonstrate the phase prediction bias caused by the filter edge effect and illustrate how forecast padding effectively addresses this issue.

#### 2) UBPP and Deadlock Rate

We first calculate the IBF and deadlock rate (*R*_*DL*_) of each algorithm as described in Supplementary Information S.7. To demonstrate the correlation between these two variables, we report Spearman’s rank correlation coefficient and its associated p-value. We present the LOWESS smoothing curves per algorithm, with 95% prediction intervals calculated using the bootstrapping method. Furthermore, assuming that negative predictions are independent events, we hypothesize that the deadlock rate can be predicted from the predictive distribution of phase using the following equation:

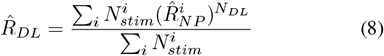

where:

- *i* is the target index (1 for 0°, 2 for 90°, etc.)
- 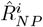 is the number of stimuli associated with target *i*
- 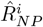 is the estimator of negative prediction probability, equal to the proportion of predicted phases outside the trigger range of target *i*
- *N*_*DL*_ is the maximum allowable number of prediction attempts per stimulus

From a graphical perspective (see Figure S7a, rightmost column, in Supplementary Information S.7), 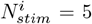 corresponds to the ratio of bin counts outside the sector associated with target *i*. In this study, *i* ranges from 1 to 4 with 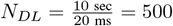 for all targets. Given that the processor makes a prediction every 20 ms and skips the trial if no target is identified within 10 seconds, 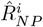.

To assess the reliability of the proposed predictor, we conducted both within-session and cross-session validations by comparing the predicted values with the actual deadlock rates in the fake-pulse sessions. In within-session validation, we predicted the deadlock rate using 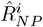 from the same session. For cross-session validation, we used 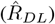 computed from the first fake-pulse session to predict the deadlock rate in the second session of the same subject. We excluded subject s18 from cross-session validation due to having only one fake-pulse session, which prevented proper partitioning into training and validation sets.

## III. Results

### A. Predictive Performance and Robustness Evaluation

As shown in Figure 3, we observed significant performance improvements across all tested algorithms, including three commonly used algorithms (AR [11], ETP [12], FFT [17]) and one newly proposed algorithm (PR). Notably, even the FFT algorithm, which previously showed the worst performance, achieved superior results with forecast padding compared to the best-performing non-forecast-padded algorithm (ETP) (right-tailed Wilcoxon signed-rank test, *p* = 3.188 *×*10^*−*14^). Beyond improved predictive performance, forecast padding significantly mitigated the UBPP in three of the four algorithms, suggesting reduced deadlock rates in real-time systems. Furthermore, as detailed in Supplementary Information S.9, the predicted signals from existing non-forecast-padded algorithms proved less informative than zeros padded beyond the signal’s end, while forecast padding substantially enhanced the quality of predicted information.

**Fig. 3:**
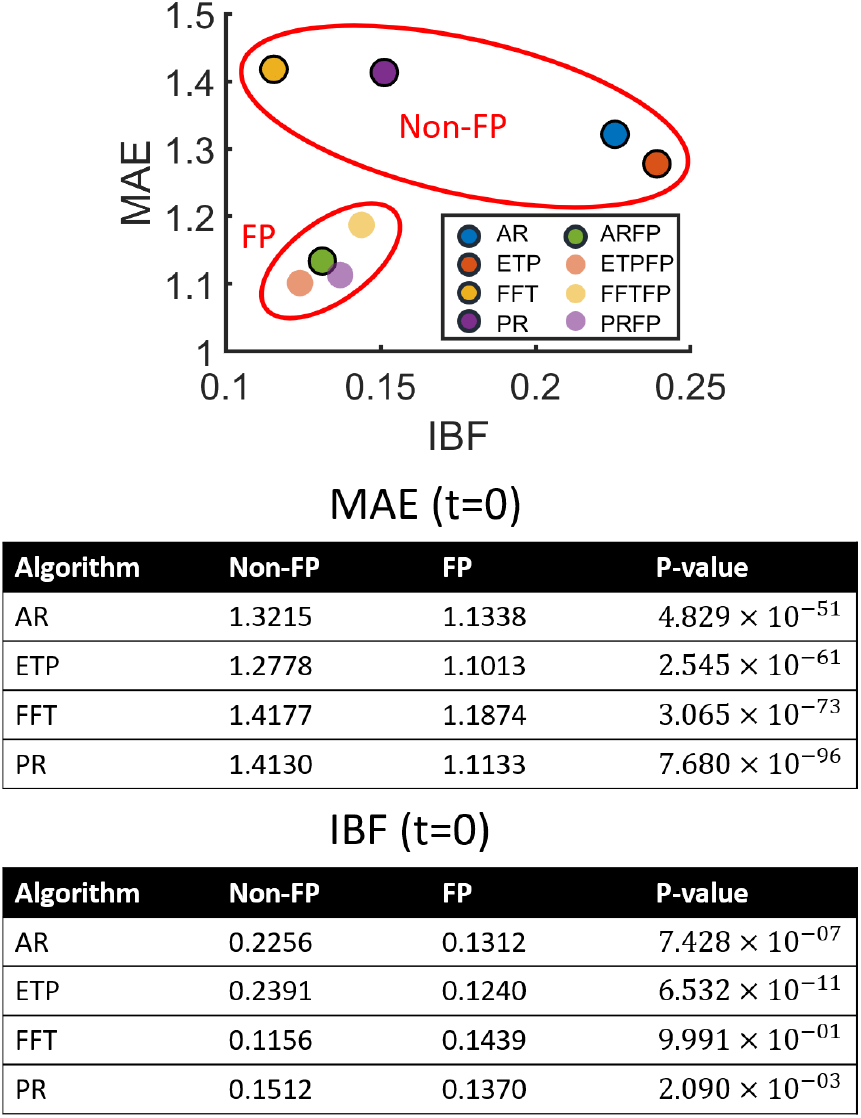
Forecast padding improves both predictive performance (MAE) and robustness (IBF) of all algorithms. Black marker edges indicate algorithms implemented in the real-time system for further investigation. Right-tailed Wilcoxon signed-rank test demonstrates significant improvement except for the IBF of FFT. MAE ranges from 0 to *π*, where 0 indicates perfect prediction and *π* indicates completely antiphasic prediction. IBF ranges from 0 to 1, where 0 indicates uniform prediction and 1 indicates purely unbalanced prediction.

**Fig. 4:**
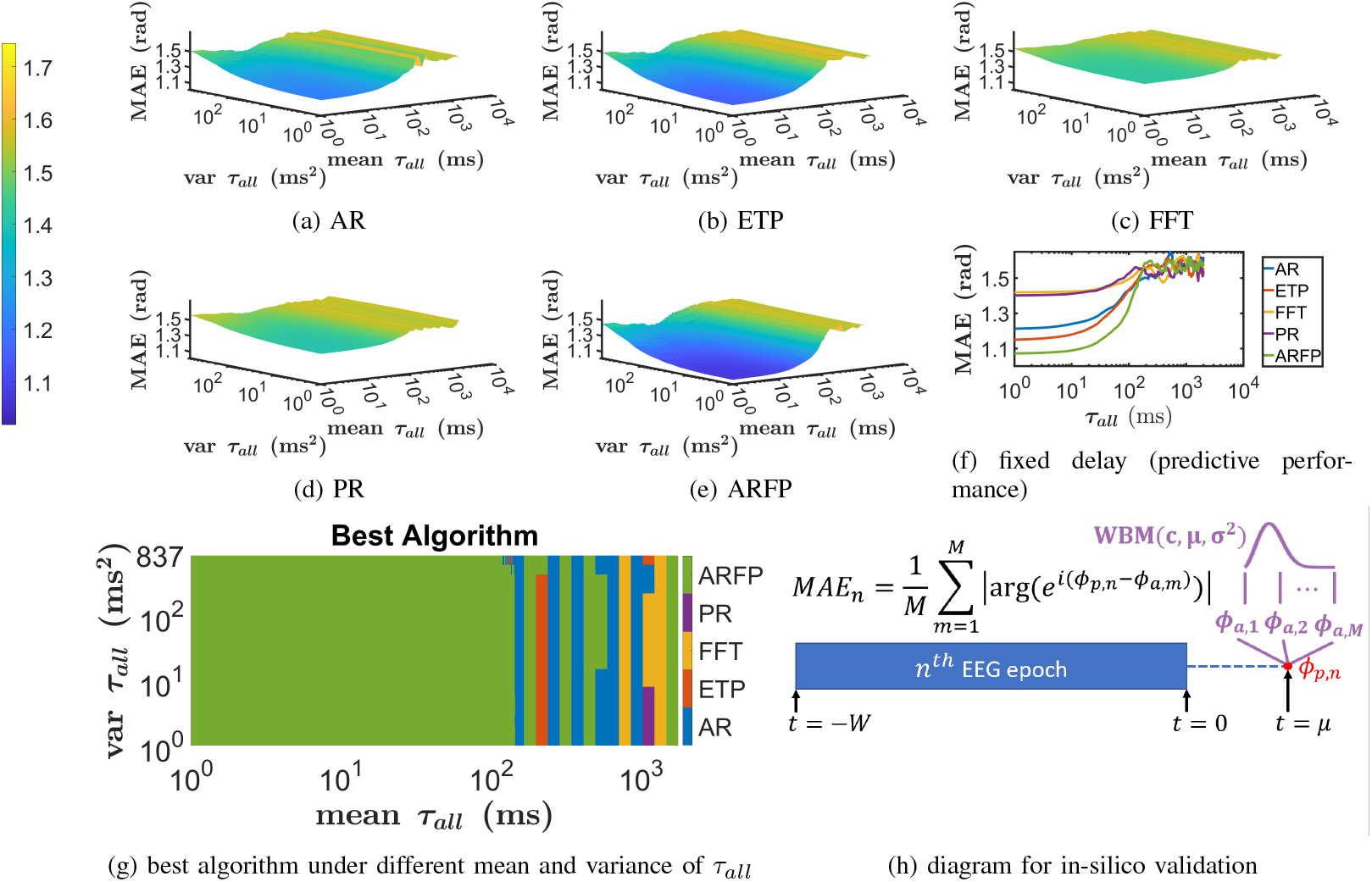
In-silico validation of stimulation accuracy. (a) to (e) MAE of each algorithm under different mean and variance of *τ*_*all*_, parameterized by *µ* and *σ*^2^ of the Weibull maximum distribution (c=279.45) respectively. The algorithm with lowest MAE for each mean and variance of *τ*_*all*_ is color-coded in (g). (f) the predictive performance under different prediction lengths. In this case, *τ*_*all*_ is assumed constant across stimuli, indicating the intersection curves of the plane *var τ*_*all*_ = 0 and the surfaces in figures (a) to (e). (h) the procedures for in-silico validation

### B. Validation Framework for Stimulation Accuracy

#### 1) In-Silico Validation

Figure 4f demonstrates the algorithms’ predictive performance across varying prediction lengths. Figures 4a through 4e illustrate the stimulation accuracy, incorporating both predictive performance and system delay jitter. As the mean or variance of *τ*_*all*_ increases, the MAE approaches *π/*2 radians, corresponding to random prediction performance. Our results demonstrate that ARFP consistently outperforms non-forecast-padded algorithms in both predictive performance and stimulation accuracy. The improvement from AR to ARFP particularly signifies the efficacy of forecast padding. As shown in Figure 4g, ARFP achieves the highest stimulation accuracy within practical ranges of *τ*_*all*_ mean and variance, demonstrating that forecast padding enables robust prediction performance even in high-latency systems. Additionally, we quantify the speed-accuracy tradeoff for each algorithm by evaluating the predictive performance at *t* = 𝔼 (*τ*_*algo*_). This arises from the fact that when *τ*_*sys*_ = 0, the prediction length is at the lower bound (i.e., 𝔼 (*τ*_*all*_) = 0 + 𝔼 (*τ*_*algo*_)), and the resulting performance cap is consequently equal to *MAE*(*t* = 𝔼 (*τ*_*algo*_)). ARFP exhibits the best speed-accuracy tradeoff among all algorithms (AR: 1.3690, ETP: 1.2650, FFT: 1.4615, PR: 1.4455, ARFP: 1.1076 radians), demonstrating that forecast padding is beneficial in low-latency systems as well.

#### 2) Fake-Pulse Validation

During the online experiment, some stimuli were skipped due to communication deadlock. MAEs are reported for both all stimuli and exclusively for non-skipped stimuli. Figure 5a shows the MAE of the five algorithms for non-skipped stimuli. ARFP demonstrates the lowest pooled MAE (incorporating all four target phases) and achieves the best performance at 180° and 270°. The runner-up, ETP, shows the second-best pooled MAE and optimal performance at 0° and 90°. The Kruskal-Wallis test reveals that the pooled MAEs of AR, ETP, and ARFP are significantly lower than those of the other two algorithms, though no significant differences exist among these three. Notably, ETP tends to skip the “difficult trials”: As shown in Figure 5b, when including MAEs from skipped stimuli, ETP’s performance ranking drops to third place for 90° and pooled MAE, while ARFP maintains its superior accuracy.

**Fig. 5:**
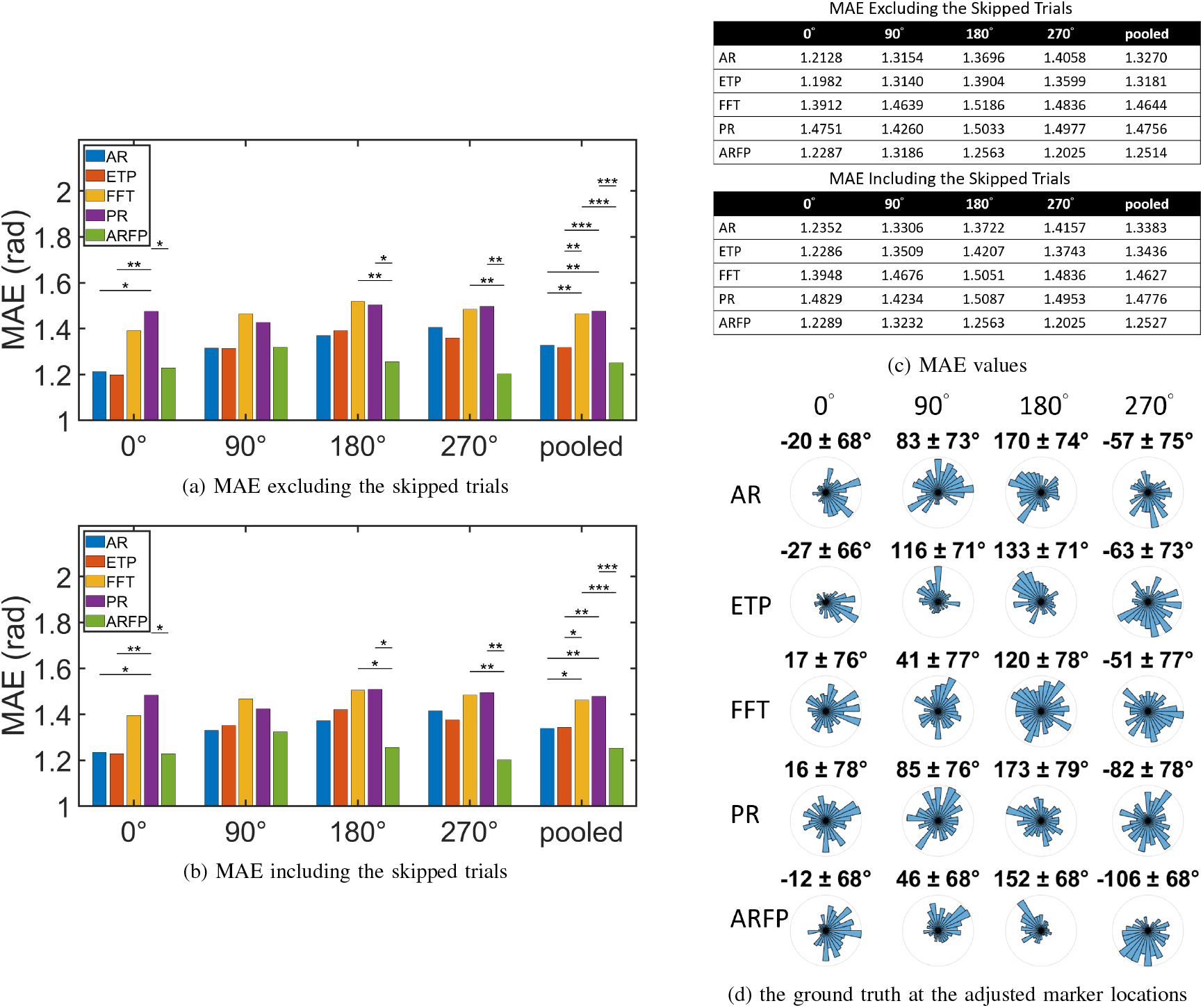
Fake-pulse validation. (a) and (b) Fake-pulse MAE for non-skipped stimuli only and for all stimuli, reported across the 5 algorithms and 5 phase conditions (4 target phases (0°, 90°, 180°, 270°) and the pooled results across all phases). Note that for skipped (forcefully terminated) stimuli, the predicted phase *ϕ*_*p*_ is not within the trigger range of the target phase. For each condition, Kruskal-Wallis test with Tukey-Kramer post-hoc test was applied to test the difference in median MAE among algorithms. Single asterisk indicates *p ≤* 0.05, double asterisks indicate *p ≤* 0.01, and triple asterisks indicate *p ≤* 0.001. (c)MAE values of each bar in figures (a) and (b). (d) The polar histograms depict the ground truth at the adjusted marker locations for each algorithm and target phase. The circular mean and circular standard deviation, presented as “mean ± standard deviation,” are denoted above each plot.

#### 3) Real-Pulse Validation

We compared the stimulation accuracy between AR and ARFP in the real-pulse session to scrutinize the effect of forecast padding. Figure 6 demonstrates that the pre-pulse waveforms for both algorithms are effectively synchronized at 0°, 90°, and 180°. However, at 270°,

**Fig. 6:**
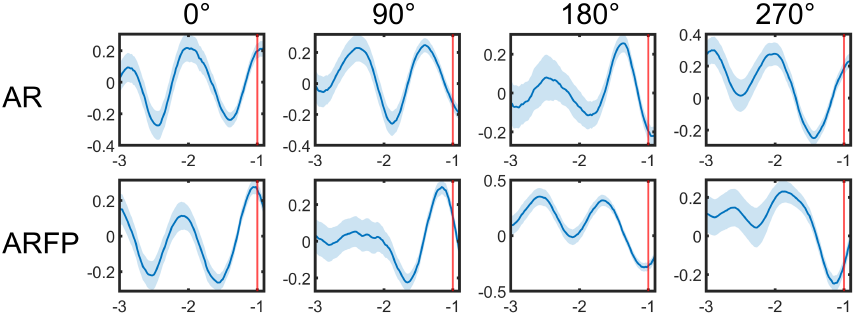
Real-pulse validation. The waveform is averaged across stimuli for each algorithm and each target phase. The time axis is normalized to the per-epoch cycle of the *µ* rhythm, denoted as *T*_*µ*_. The blue curve and the shaded region indicates the mean ± standard error. The red vertical line indicates one *T*_*µ*_ before the pulse onset.

AR’s waveform appears to target the peak (0°) rather than the intended equilibrium point (270°), consistent with its higher MAE observed in fake-pulse validation. In contrast, ARFP maintains satisfactory performance at 270°, further confirming its superior phase-targeting capability across all target phases.

### C) Validation Framework for Robustness

#### 1) Filter Edge Effect and UBPP

The intermediate steps of AR and ARFP are demonstrated in Figure 7. The benchmark signal in the left column demonstrates that AR and ARFP share similarly distributed ground truth, ensuring that the different predictive distribution for these two algorithms is not due to the underlying ground truth differences. The middle column depicts the resulting signal from Step 1. Due to the zero padding, both algorithms suffer from the vanishing amplitude at the end. This vanishing trend inclines the autoregressive model to predict a very small amplitude around t=0, resembling the equilibrium points (90° and 270°) of a cosine signal. The right column reveals the resulting signal from Step 2. Benefiting from the forecast padding, the waveform of ARFP at the right end is diversified, compared to the severely diminished amplitude of AR near t=0. This explains the more abundant predicted current phase of ARFP compared to AR.

**Fig. 7:**
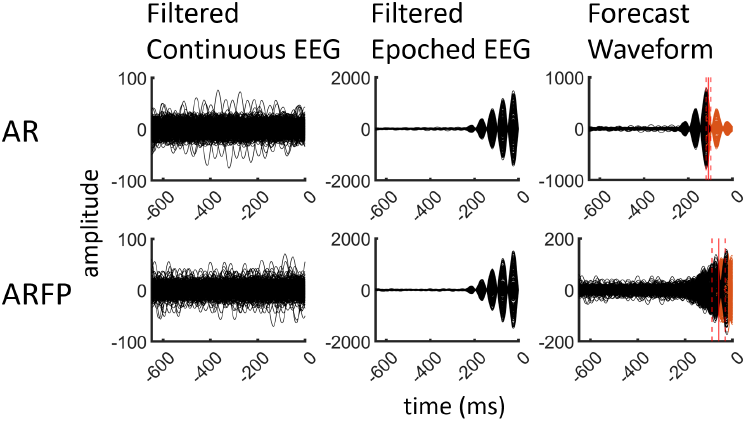
Intermediate steps of AR and ARFP from 1000 EEG epochs. The left column shows the benchmark *µ* rhythm obtained from continuous EEG filtering. The middle column displays the resulting signal from the first step (zero-phase bandpass filtering). The right column exhibits the resulting signal from the second step, where the orange curves indicate the time intervals of forecast portion. The vertical solid red line and the dashed line aside indicate the mean ± standard deviation of the length of the removed edge portion of signals.

#### 2) UBPP and Deadlock Rate

Figure S7 in Supplementary Information S.7 reports the IBF and deadlock rate for each algorithm. ARFP, enhanced by forecast padding, demonstrates the lowest IBF and deadlock rate. Figure 8a reveals strong to moderate correlations between IBF and deadlock rate across all algorithms. Furthermore, Figure 8b validates the high precision and accuracy of the proposed predictor in both within-and cross-session validation. While the within-session validation confirms the relationship between UBPP and deadlock rate, the cross-session validation demonstrates the predictor’s high generalizability, suggesting that an algorithm’s predictive distribution on prerecorded resting-state EEG can effectively predict the deadlock rate in an actual system.

**Fig. 8:**
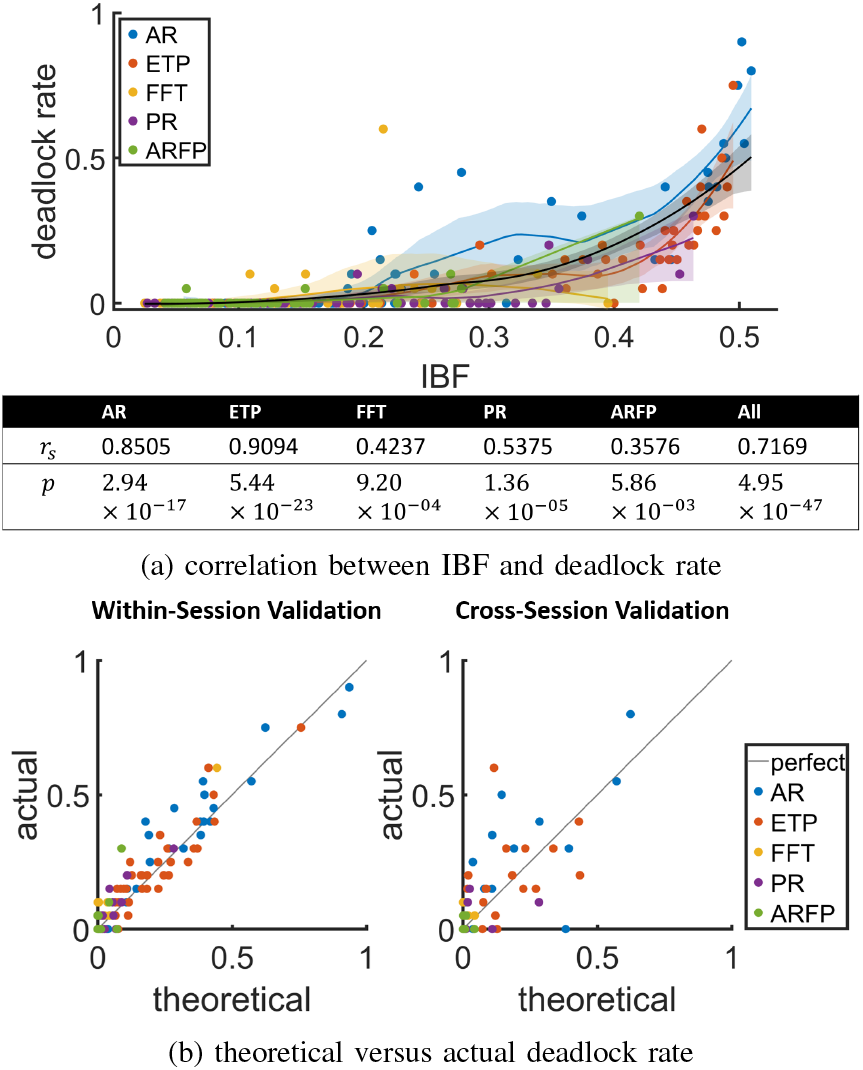
Robustness indicator and predictor. (a) Correlation between IBF and pooled deadlock rate. Each data point corresponds to a fake-pulse session. The deadlock rate is calculated for each algorithm by dividing the number of skipped trials by the overall number of stimuli. The IBF is derived from all prediction attempts for the algorithm, not solely those involving stimulus presentation. The solid line and the shaded region around it represent the LOWESS smoothing curve and the 95% prediction interval for each algorithm, as well as for the aggregate of all algorithms (depicted by the black curve and shading). The table below the figure provides the Spearman’s rank correlation coefficient and the associated p-value. (b) Theoretical 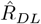 versus actual (*R*_*DL*_) deadlock rate. The gray line indicates a perfect prediction. For within-session validation, each point corresponds to a fake-pulse session, while for cross-session validation, each point corresponds to a subject. The root-mean-square error (RMSE) between 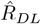 and the actual deadlock rate *R*_*DL*_ in the within-session validation for each algorithm is AR: 0.0572, ETP: 0.0495, FFT: 0.0301, PR: 0.0248, ARFP: 0.0329, and the RMSE in the cross-session validation is AR:0.1558, ETP:0.1508, FFT:0.0346, PR:0.0604, ARFP:0.0175.

## IV. Discussion

### A. Forecast Padding in Personalized Neurostimulation

Forecast padding has potential applications in closed-loop therapies for various neurological disorders. We demonstrate two key application scenarios in neurostimulation where forecast padding shows particular promise.

#### 1) Responsive Neurostimulation for Seizure Suppression

Responsive neurostimulation (RNS) is a treatment that monitors brain activity and delivers electrical stimulation to specific areas of the brain when it detects patterns associated with an impending seizure. In this approach, early detection and intervention are crucial to minimize the impact of seizures and maximize the therapeutic window [28], [29], [30]. Current RNS systems typically operate with detection latencies ranging from 2 to 12 seconds [28], indicating significant room for improvement. Forecast padding’s ability to preserve boundary information could potentially reduce these latencies by providing more accurate signal representations at processing boundaries, thereby enabling faster and more reliable seizure detection. The improved signal fidelity at signal boundaries might also increase the accuracy of seizure detection, although this aspect requires further investigation.

#### 2) Closed-Loop TMS for Depression Treatment

Recent studies have been exploring the potential of EEG-phase-synchronized TMS as a personalized treatment for depression [9], [7]. However, the overly long inter-pulse interval due to system deadlock [14], [7], [15] may obstruct the development of effective treatment protocols. Specifically, inter-pulse/burst intervals in rTMS [31], [32] and TBS [33], [34] play a crucial role in modulating synaptic efficiency; Overly low pulse rate, therefore, may not result in intended long-term potentiation/depression. We propose that forecast padding, by reducing deadlock occurrence and enhancing overall accuracy through improved boundary processing, is essential for developing clinically viable phase-synchronized rTMS or TBS protocols. With more precise timing control in stimulation protocols and appropriate inter-pulse/burst intervals, phase-synchronized TMS with forecast padding may improve therapeutic outcomes in neurological and psychiatric disorders.

### B. Forecast Padding in Other Fields

While our primary focus has been on medical applications, forecast padding has broader potential to benefit various signal processing domains, as the kernel is adaptable to meet specific processing requirements. A notable application lies in convolutional neural networks (CNNs). In most CNN models, zero-padding is employed to maintain spatial dimensions [35]. However, zero-padding causes boundary effects and compromises the translation invariance property of CNN [36], [37], thereby degrading model performance across various tasks. To address these limitations, recent approaches have utilized image border statistics for padding generation, demonstrating superior performance to zero padding in various computer vision tasks [38], [39], [40]. Compared to these methods, forecast padding’s unique approach of utilizing kerneldistilled information for padding generation enables task-specific optimizations. Future work may evaluate this kernel-aware padding strategy across different CNN architectures and applications, such as real-time object detection in autonomous driving or high-frequency trading in finance, where boundary effects significantly impact performance.

### C. Limitations and Future Directions

Despite these promising applications, several important considerations and future directions warrant discussion. First, exploring forecast padding with hybrid approaches that combine different algorithms (e.g., using the predicted signal from FFT to pad the input signal for AR models) could leverage complementary strengths for higher prediction accuracy, particularly in scenarios where individual algorithms show distinct advantages under varying signal conditions.

Second, our current validation focused on task-free EEG; future studies should examine forecast padding’s performance under more dynamic, task-based conditions to assess its generalizability across different brain states. This extension is particularly relevant for clinical applications where patients may be engaged in cognitive tasks or experiencing varying levels of arousal during treatment.

Finally, our proposed predictor of deadlock rate assumes that negative predictions are independent events. More sophisticated probabilistic approaches, such as Bayesian inference or Markov modeling, could better account for potential systematic phase drifts and correlated prediction errors. These methods could provide more nuanced estimates of deadlock probabilities and lead to adaptive strategies that optimize stimulation parameters to balance accuracy and deadlock rate based on real-time signal characteristics.

Collectively, addressing these limitations would advance the robustness and clinical utility of phase-synchronized TMS systems across various experimental paradigms and therapeutic interventions.

## V. Conclusion

Effective and personalized closed-loop neuromodulation requires interdisciplinary collaboration among neuroscientists, clinicians, and engineers. This paper addresses technical challenges arising from causality limitations, paving the way for more sophisticated therapeutic protocols and devices. Specifically, we use a novel delay-relevant framework to demonstrate the significant improvements in accuracy and robustness achieved through forecast padding. We hope this technical breakthrough will contribute to the advancement of closedloop neuroscience.

