## Supplementary Information for "Forecast Padding Enhances Accuracy and Robustness of EEG-Phase-Synchronized TMS"

### Preliminary

*Abstract:* This preliminary article introduces key concepts of EEG-phase-synchronized TMS, laying the groundwork for a deeper understanding of the main article.

#### 1. Delay Model of Closed-Loop TMS

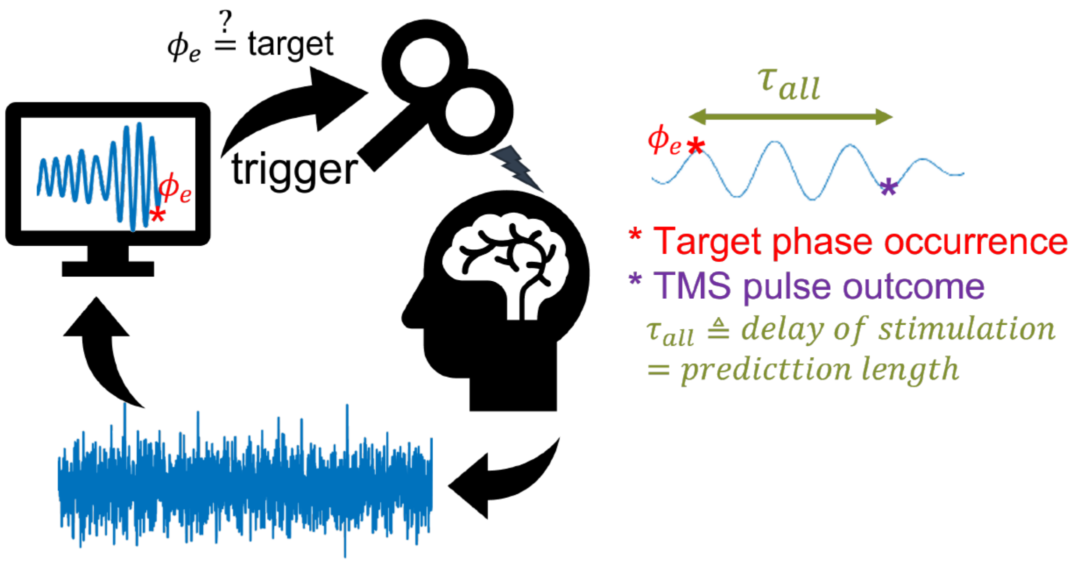


Figure P1: Diagram of EEG-phase-synchronized TMS. A stimulus is delivered when the predefined neural biomarker (e.g., phase of the EEG signal) is detected. To ensure the stimulus occurs precisely at the target phase, the transport delay must be compensated through prediction. This compensation is achieved by setting the prediction length equal to the transport delay.

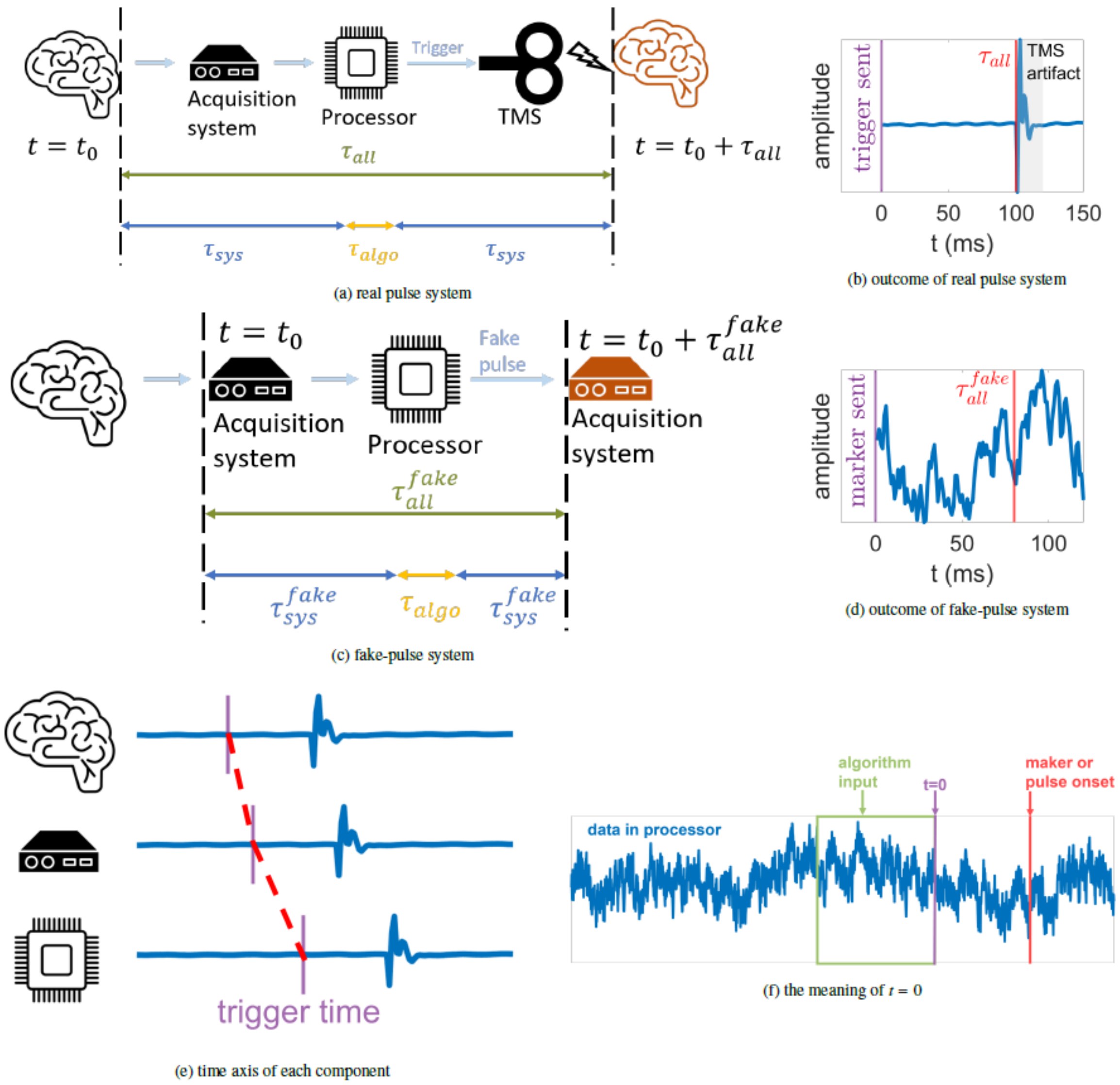


Figure P2: Illustration of the delay model proposed in [4]. (a) and (b) System diagram in (a) is adapted from our prior work [4]. A closed-loop TMS system consists of an acquisition system, a processor, and a TMS stimulator. The transport delay, denoted as $\tau_{all}$, can be measured from the interval between the trigger time ($t = 0$) and the onset of the TMS pulse (see Supplementary S.1 for the derivation). $\tau_{all}$ can be decomposed into algorithmic delay ($\tau_{algo}$) and non-algorithmic delay ($\tau_{sys}$). The non-algorithmic delay $\tau_{sys}$ encompasses processing, queueing, transmission, and propagation delays within the closed-loop network, while the algorithmic delay $\tau_{algo}$ stems from the computational time of the phase prediction algorithm. (c) and (d) To avoid TMS artifact blurring the EEG signal and disable the calculation of pulse phase, the fake-pulse system is derived from the real-pulse system by replacing the TMS trigger with an event marker (fake pulse). Substituting markers for TMS triggers allows for acausal estimation of the ground truth phase. The latency between marker transmission and reception is denoted as $\tau_{all}^{fake}$ and can also be divided into algorithmic ($\tau_{algo}$) and non-algorithmic ($\tau_{sys}^{fake}$) components. This substitution modifies the non-algorithmic delay from $\tau_{sys}$ to $\tau_{sys}^{fake}$, while leaving the algorithmic delay $\tau_{algo}$ unaffected. (e) and (f) The delay introduces temporal misalignment among various system components, and the red dashed line connects corresponding data samples on each time axis. When quantifying $\tau_{all}$ and $\tau_{all}^{fake}$, the starting and ending points must be aligned on the same time axis. In this study, the processor's time axis is employed, and the current time ($t = 0$) has three equivalent meanings - the most recent EEG data point received by the processor, the final point of the EEG epoch input to the algorithms, the trigger time of the event marker or TMS pulse.

In a phase-synchronized TMS system, stimuli are delivered according to the phase of the ongoing EEG rhythm. The meaning of EEG phase will be detailed in Preliminary Section 2, and readers can currently consider it as the biomarker to determine the timing of stimuli. In some studies, the processor continuously updates its data window and uses the latest window to calculate the phase at the current time (t=0). TMS is triggered when the current phase matches a predefined target phase [2, 3]. However, this strategy is effective only when the system's delay is negligible. In practical closed-loop TMS systems, there exists a latency denoted as $\tau_{all}$ between the trigger command and the actual TMS pulse outcome, as shown in Figure P1. When the TMS pulse is triggered, the estimated target phase at the current time point has already elapsed. The origin of the delay is described in [4] and is briefly summarized in Figure P2. To account for $\tau_{all}$, two trigger strategies are commonly employed:

**Strategy 1. Trigger-Range Strategy:** This strategy involves utilizing the most recent $\mathrm{W}$ samples (from $\mathrm{t}=-\mathrm{W}+1$ to $\mathrm{t}=0$) received by the processor and predicting the phase at $\tau_{all}$. If the predicted phase falls within the predefined trigger range, the TMS is triggered. If not, the algorithm waits for additional incoming data and uses the most recent $\mathrm{W}$ samples to make a prediction again. The above process repeats until a positive prediction occurs (i.e., the target is predicted to occur at $\tau_{all}$). The overlapping portion of the EEG data window between consecutive prediction attempts depends on the processing rate of the algorithm.

**Strategy 2. Timer Strategy:** In this approach, the algorithm employs the most recent $\mathrm{W}$ samples (from $\mathrm{t}=-\mathrm{W}+1$ to $\mathrm{t}=0$) to predict the future time of the upcoming target, denoted as $\tau_{target}$, which is at least $\tau_{all}$ away from now (i.e., $\tau_{target}>\tau_{all}$). Subsequently, a delay timer is set to trigger the TMS at the time $\mathrm{t}\ =\ \tau_{timer}=\tau_{target}-\tau_{all}$ from the present moment.

Since both strategies involve prediction of future phases and the prediction horizon increases as $\tau_{all}$ grows, it is intuitive to expect a decrease in prediction accuracy as $\tau_{all}$ increases. However, many studies have disregarded the impact of this delay on prediction performance, leading to overestimated stimulation accuracy in their validation results. Preliminary Section 3 will describe in detail how the stimulation accuracy is overestimated, but before that, we will introduce how the phase of the EEG rhythm is computed.

#### 2. Phase Prediction Algorithm

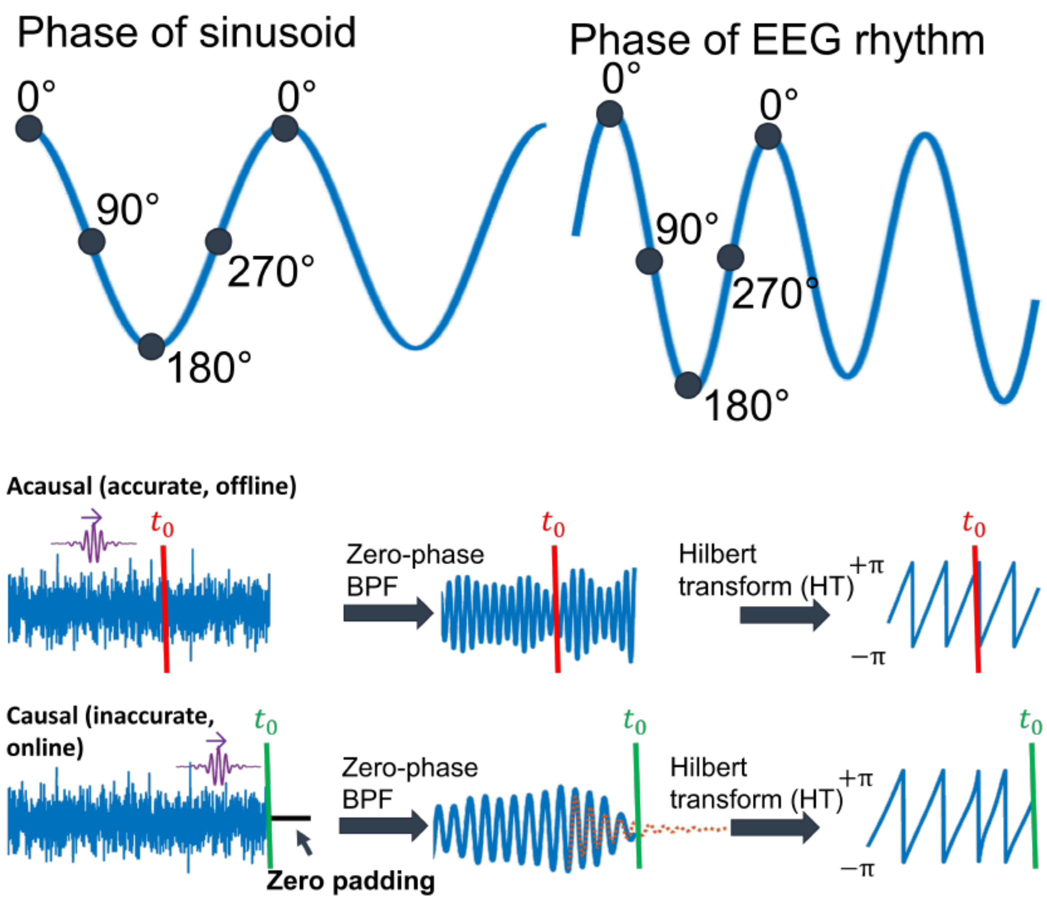


Figure P3: Calculation of EEG phase.

The phase of a sinusoidal signal indicates the position within its waveform cycle. For instance, a phase value of 0° corresponds to the peak, 180° to the trough, and 90°/270° represent the midpoints of the falling/rising edge (see Figure P3). EEG phase definition leverages the phase-waveform connection as seen in sinusoids. A common definition involves calculating the argument of the analytic signal [7]. Formally, let $\mathrm{x(t)}$ represent the raw EEG signal. The EEG rhythm $\mathrm{x}_b(t)$ can be estimated by zero-phase bandpass filtering $\mathrm{x(t)}$ to extract the desired frequency band. Next, the bandpass signal $\mathrm{x}_b(t)$ is converted to the analytic signal $\mathrm{x}_a(t)$ using:

$$\mathrm{x}_a(t) = \mathrm{x}_b(t) + i\mathrm{x}_h(t)$$

, where $\mathrm{x}_h(t)$ is the Hilbert transform of $\mathrm{x}_b(t)$.

The phase of $\mathrm{x}_b(t)$ is defined as the argument of the analytic signal:

$$\phi(\mathrm{t}) := \mathrm{atan2}(\mathrm{x}_h(t), \mathrm{x}_b(t))$$

, where $\mathrm{atan2}$ denotes the 2-argument arctangent. As depicted in Figure P3, the 0° and 180° of $\phi(\mathrm{t})$ correspond to the EEG rhythm's peak and trough, respectively, with 90° and 270° delineating the falling and rising edge midpoints. This method is known as acausal phase estimation because zero-phase filtering uses past and future signal to determine the output at a given time (In signal processing, "acausal" refers to a system or signal where the output at any given time depends on both past and future inputs.) Despite proven reliable in extracting the physical oscillatory phase and frequency [7-9], the acausal phase estimation is unsuitable for closed-loop TMS due to its reliance on future signals, unavailable in real-time scenarios.

To address this limitation, various causal phase estimation algorithms have been proposed [1, 10-12]. These algorithms estimate the phase at the current time using only preceding data, making them well-suited for real-time closed-loop TMS applications.

Since future data is unavailable during filtering, these approaches either:

- Apply causal filters to the signal [12].
- Pad zeros beyond the signal's end before zero-phase filtering [10, 11].

In both cases, prediction of future phases is inevitable due to the group delay of the causal filter or the data distortion around the edge. For example, with zero-phase filtering, algorithms might:

1. Pad zeros beyond the signal's end.
2. Apply zero-phase bandpass filtering to the padded signal.
3. Remove distorted edge portions due to filter effects.
4. Use the remaining data to predict future phase or signal (dashed red line in Figure P3). If a signal is predicted rather than phase (e.g., autoregressive phase estimation proposed by Chen et al. [10]), the Hilbert transform is employed to convert the predicted signal into the instantaneous phase.

3. Overrated Stimulation Accuracy and Irreproducible Neurological Findings

| Paper | Validation Type | Trigger Strategy | Performance Metric | Issues |
|---|---|---|---|---|
| Chen et al., 2011 [10] | IS | TIM | Bias/var at $t = \tau_{target}$ | Issue 1 (in IS) |
| Zrenner et al., 2018 [2] | FP/RP | TRIG | Bias/var at $t = 0$; waveform preceding TMS pulse | Issue 1 (in FP) |
| Shirinpour et al., 2020 [11] | IS/FP | TRIG | Bias/var/MAE-based accuracy at $t = 0$ | Issue 1 (in FP) |
| Mcintosh & Sajda, 2020 [1] | IS | NPS | MAE at $t = 0$ | Issue 1 (in IS) |
| Mansouri et al., 2017 [12] | IS | NPS | PLV between prediction and ground truth across $\tau_{all}$ | Issue 3 (in IS) |
| Schatza et al., 2022 [13] | IS/FP | TRIG | Bias/var at $t = \tau_{all}^{fake}$ | Issue 2 (in FP) |

| | | | | |
|---|---|---|---|---|
| Madsen et al., 2019 [3] | FP/RP | TRIG | Bias/var/MAE at $t = 0$; waveform preceding $t = 0$ | Issue 1 (in FP/RP) |
| Faller et al., 2019 [14] | FP/RP | TIM | PPV; phase distance between target and markers | Issue 2 (in FP) |
| Tseng et al., 2023 [15] | IS/FP/RP | TRIG | Phase distance between target and stimuli | Issue 2 (in FP) |

Table P1: Comparison of the existing validation methods. IS: in-silico validation; FP: fake-pulse validation; RP: real-pulse validation; TRIG: trigger-range strategy; TIM: timer strategy; NPS: non-phase-synchronized, the validation does not target a specific phase, but simply compares the causally predicted phase to the ground truth; bias: the mean circular distance between the target phase and the pulse phase (ground truth at the actual or simulated pulse locations) ; var: abbreviation for "variance", the spread of the pulse phase around its circular mean; MAE: mean absolute error (also called MACE in [1]) between the predicted phase and the ground truth; PLV: phase-locking value, ranging from 0 to 1, representing no to perfect phase synchronization. PPV: positive predictive value, ratio of pulses falling in the correct half-cycle.

Although impractical for real-time system, the acausal phase estimation is typically used as a ground truth (GT) to evaluate the performance of the causal algorithms and the accuracy of phase-synchronized stimuli [1, 2, 11, 12]. Specifically, the phase error is computed as the difference between the causally predicted target phases and the ground truth phases at the time of TMS pulse onset (referred to as pulse phase hereafter). The error can be decomposed into two primary components: bias and variance [10, 11]. Bias represents the offset between the target phase and the central point of the pulse phase distribution, indicating how effectively the system locks onto the target phase. In contrast, variance quantifies the degree of dispersion in the pulse phase distribution. In this study, mean absolute error (MAE) is selected as the accuracy metric [1, 3] because it incorporates the influence of both bias and variance. MAE is defined as $\mathrm{MAE} = \frac{1}{N}\sum_{n=1}^{N}\left|\arg\left(e^{i(\phi_{p,n}-\phi_{a,n})}\right)\right|$, where $N$ is the number of prediction trials, $\phi_{p,n}$ is the predicted phase of the causal algorithm, $\phi_{a,n}$ is the ground truth at the pulse onset, and $\arg$ is the wrapped phase of the complex number between $-\pi$ and $\pi$. The range of MAE from $0$ to $\pi$ indicates the accuracy of the prediction, ranging from a perfect prediction to a completely anti-phasic prediction.

One common challenge in assessing MAE arises from the presence of TMS artifacts in the EEG data. This impedes the application of the acausal approach to extract the ground truth. To address this issue, several validation approaches have been proposed.

##### 3.1 Fake-Pulse Validation

The first approach, referred to as fake-pulse validation, is to employ non-stimulating trials for ground truth evaluation (see Figure P2c and Figure P2d). This approach involves triggering event markers in a phase-synchronized way and storing trigger events within the EEG data to mark the timing of the TMS delivery even though no actual TMS pulse was delivered [2, 3, 11, 13, 14]. The MAE is then derived between the phases predicted in real time and the ground truth at the event locations.

##### 3.2 In-Silico Validation

The second approach is referred to as in-silico validation. To mimic the real-time scenario, this approach reads in pre-recorded resting-state EEG (rsEEG) data and simulates phase-synchronized stimuli [1, 10-13]. In this case, the ground truth at the simulated pulse locations is available because there is no actual TMS artifact, enabling the comparison of predicted phases to the ground truth.

##### 3.3 Real-Pulse Validation

The third approach, referred to as real-pulse validation, involves delivering actual TMS in a phase-synchronized way and evaluate the accuracy using a quantitative or qualitative analysis. The quantitative analysis involves reconstructing the post-TMS signal and subsequently calculating the ground truth using an acausal method [11, 14]. However, it remains uncertain whether the reconstruction process introduces systematic errors in phase estimation. Even in cases of perfect reconstruction, acausal phase estimation may not have any physical meaning due to the TMS-induced phase reset in brain rhythms [16, 17], causing phase discontinuities at the onset of the TMS pulse. On the contrary, qualitative analysis simply plots the average waveform before pulse onset to visualize whether the waveform resembles the target phase (e.g., Figure 4d in [2]).

##### 3.4 Issues in Validation

Despite the diversity of validation approaches, three primary issues persist in existing studies:

**Issue 1. ignoring $\tau_{all}$ in real-pulse, fake-pulse, and in-silico validation:**

This issue indicates that the accuracy is validated at $\mathrm{t} = 0$ instead of $\mathrm{t} = \tau_{all}$. This insufficient consideration is associated with various validation approaches and trigger strategies. For real-pulse validation, an error example is to plot the averaged waveform time-locked to the trigger time instead of the pulse onset [3]. For fake-pulse validation, an error example is to compute the error between ground truth and the causally estimated phase at the trigger time ($\mathrm{t} = 0$), instead of $\mathrm{t} = \tau_{all}$ [2, 3, 11]. For in-silico validation, an error example is to compute the error between the causally and the acausally estimated phase at $\mathrm{t} = 0$ [1, 11]. Another error example with timer strategy is to omit the constraint $\tau_{target} > \tau_{all}$ by setting $\tau_{all}$ to zero [10]. For all the examples listed above, the ignoring of $\tau_{all}$ led to inflated accuracy in their validation results.

**Issue 2. Assuming same delay for marker and TMS pulse in fake-pulse validation:**

This faulty validation ignores the difference between the transport delay of a fake-pulse system and a real-pulse system, implicitly expecting TMS pulses ($t = \tau_{all}$) to occur at the marker location ($t = \tau_{all}^{fake}$) [13, 14]. Specifically, the researchers compared the causally predicted phases to the ground truth at the marker locations. However, since $\tau_{all}$ is usually greater than $\tau_{all}^{fake}$ in practical situation, this assumption can lead to an underrated prediction horizon and overrated accuracy.

**Issue 3. Ignoring jitter of $\boldsymbol{\tau_{all}}$:**

Many studies reported the predictive performance as a function of prediction length [12, 18], particularly when in-silico validation is conducted. As a result, readers can easily misinterpret an algorithm's predictive performance as stimulation accuracy. In fact, the latter is affected by both the former and the jitter of $\tau_{all}$ in the real system. Specifically, during the online setting, the phase is predicted at a fixed time point, $t = \mathbb{E}(\tau_{all})$, where the expectation value is calculated as the empirical mean of the past data. However, the actual pulse onset varies from trial to trial, with the variance denoted by $\mathbb{V}(\tau_{all})$. Given a fixed predictive performance, the stimulation accuracy decreases as $\mathbb{V}(\tau_{all})$ increases. Therefore, a valid validation framework of stimulation accuracy should simultaneously consider the predictive performance of an algorithm and the jitter of the system's delay.

The challenges and issues of existing validation methods are summarized in Table P1. Discrepancies and incomplete validation methods have led to difficulties in obtaining reliable and comparable experimental results. For example, Zrenner and colleagues [2] reported that the pre-stimulus phase of $\mu$ rhythm influences the strength of motor-evoked potentials (MEP). However, Madsen and colleagues [3] could not replicate the same phase-dependent modulation. The inconsistencies between these two studies may certainly be ascribed to the individual differences in neural processes, but another possibility is that both teams were actually targeting different phases. Specifically, both teams rely on the current phase ($t = 0$) being within the trigger range to decide whether to trigger TMS. Moreover, both teams present validation results without considering the delay (i.e., Issue 1 as mentioned above). This raises doubts that the inconsistent phase-dependent modulation effect stem from different mean pulse phases due to different $\tau_{all}$ for both systems.

4. Robustness of Algorithms

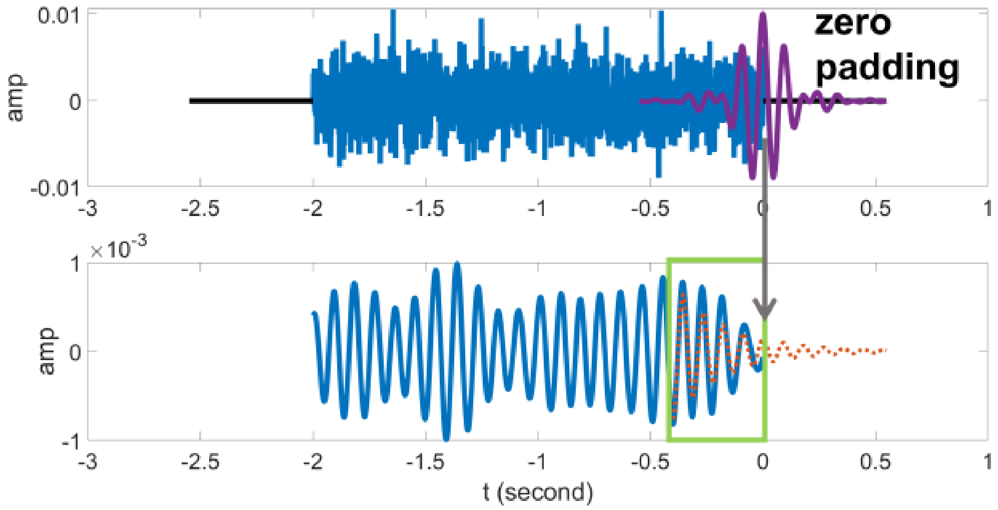


(a) Filter edge effect leads to unbalanced phase prediction.

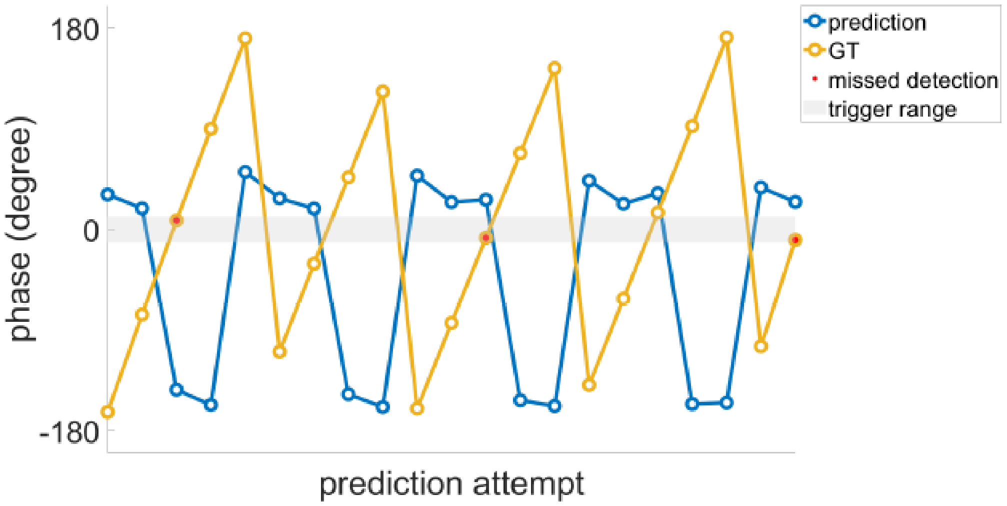


(b) Unbalanced phase prediction leads to communication deadlock.

Figure P4: Filter edge effect and communication deadlock. (a) Autoregressive phase prediction is vulnerable to the filter edge effect. The green box indicates samples removed at the right end due to distortion, and the red dashed line represents the predicted waveform. (b) The predicted phase and the corresponding ground truth are shown in one plot, illustrating the cause of communication deadlock. Here, the deadlock [5, 6] refers to the condition where the TMS machine awaits a trigger from the processor, while simultaneously, the processor waits for the completion of TMS delivery.

In addition to the performance, the unbalanced phase prediction (UBPP) is another issue that should be considered, as it can impair the robustness of the system. Using the autoregressive phase estimation approach [2] as an example, the cause and consequence of the UBPP are demonstrated in Figure P4. As shown, the first step of the algorithm is to apply zero-phase bandpass filtering to the finite-length signal. To maintain the same length after filtering, zero values are often padded beyond both ends of the signal. Let $N$ be the length of the zero-phase FIR filter (Note that if the zero-phase filtering is achived by foreward-backward filtering using a causal filter of lengh $N_1$, $N$ is equal to $2N_1 - 1$.) The $\frac{N-1}{2}$ points at both ends of the filtered signal are distorted due to the filter edge effect (Figure P4a). Therefore, $M$ points at both ends are removed and the remaining part is used to pretict the future waveform and phase up to $t = \tau_{all}$. In most cases, $M$ is chosen to be slightly smaller than $\frac{N-1}{2}$ [10, 11, 19], subject to the trade-off between the prediction length and the number fo distored data points used to fit the autoregressive model. Since the EEG samples affected by the filter edge effect are not completely removed, the autoregressive model tends to identify the vanishing trend near the end and predict a small amplitude at $t = 0$ (red dashed line), resembling the 90° and 270° points of a cosine signal. Consequently, the estimated phase at $t = 0$ is biased towards 90° and 270°, and the predicted phase at $t = \tau_{all}$ is biased towards 90°+$\Delta\theta$ and 270°+$\Delta\theta$, where $\Delta\theta$ is the phase offset associated with the $\tau_{all}$ (see Equation S10-1 in Supplementary S.10). The impact of UBPP depends on the trigger strategy. If the trigger-range strategy is used, as shown in Figure P4b, UBPP can lead to communication deadlock due to too many false negative predictions (red dots). Specifically, since the predicted phases (blue circles) are concentrated at two values, the algorithm requires numerous trials to successfully identify a target, consequently resulting in an excessively long inter-pulse interval (IPI), known as a deadlock. A long IPI impedes many clinical applications, such as phase-synchronized rTMS/TBS, as these interventions require a sufficient pulse rate to achieve therapeutic effects. In contrast, the timer strategy can resolve the deadlock problem by predicting the time of the future target, ensuring a successful trigger with each prediction. However, this approach comes at the cost of accuracy. Specifically, the unbalanced prediction, resulting in numerically close trigger time points, leads to poor stimulation accuracy. Although using the autoregressive approach as an example, it is important to note that the filter edge effect and UBPP are not limited to the autoregressive approach. Indeed, this issue can affect a wide range of causal phase prediction algorithms, provided that the computational procedure entails the use of zero-phase filtering on signals of finite length [2, 10, 11].

Supplementary Information

### S.1 Delay Measurement

Transport Delay $\tau_{all}$ Manifests Itself as the Interval between Trigger Time and Pulse Onset

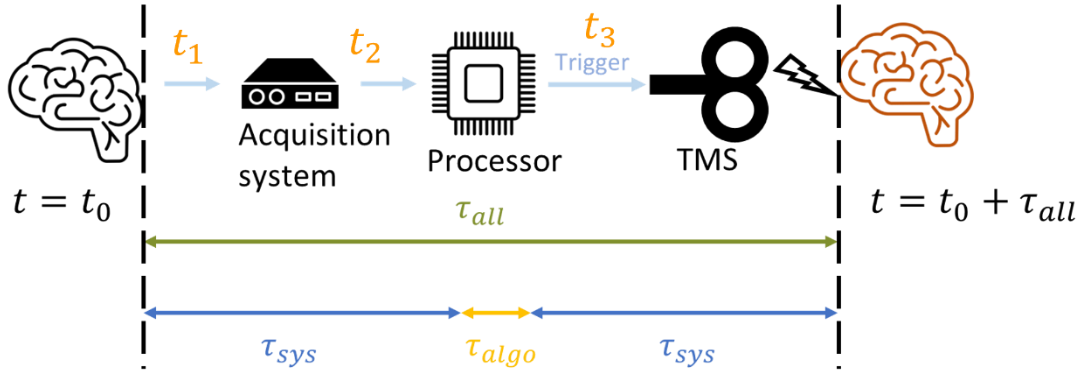


As illustrated in the above figure, $t_1$ represents the temporal discrepancy between the appearance of a specific signal point in the scalp electroencephalogram and the point at which it is detected by the acquisition system. $t_2$ signifies the time interval from when the acquisition system perceives this signal point to when it is observed by the processor. $t_3$ denotes the time lapse from when the processor detects this signal point to the occurrence of TMS pulses in the scalp electroencephalogram. The transport delay $\tau_{all}$ is equal to $t_1 + t_2 + t_3$.

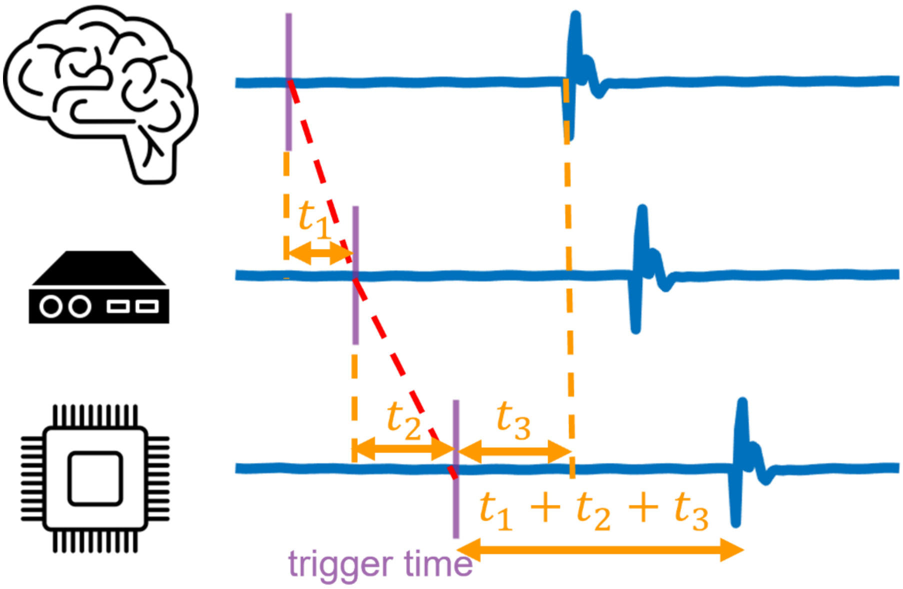

In terms of the observed EEG signal (see above figure), $t_1$ and $t_2$ corresponds to the temporal shift among the signal observed by different nodes, and $t_3$ corresponds to the lapse from when processor detect the signal point and send the trigger to the pulse onset in scalp EEG. The interval between the trigger time and pulse onset is equal to $t_1 + t_2 + t_3 = \tau_{all}$. A similar derivation can be done to show that the transport delay ($\tau_{all}^{fake}$) of the fake-pulse system is equal to the interval between the trigger time and the marker location.

Measurement of $\tau_{algo}, \tau_{all}, \tau_{sys}$ in the Real-Pulse System

The procedure for measuring $\tau_{algo}, \tau_{all},$ and $\tau_{sys}$ is as follows:

1. In the real-pulse system, the processor continuously receives EEG data and predicts future phases upon the arrival of each new data chunk.
2. For each prediction, record the computational time of the algorithm, $\tau_{algo}$, using the *tic* and *toc* functions in MATLAB.
3. When a target is detected, record the index of the current EEG sample (t=0) in the processor and trigger the TMS.
4. Denote the TMS pulse onset time for each stimulation trial in the offline analysis.
5. $\tau_{all}$ is calculated as the interval between the recorded trigger time and the subsequent pulse onset.
6. $\tau_{sys}$ is calculated as $\tau_{all}$ minus $\tau_{algo}$.

The statistics below show the mean ± standard deviation of delay values over stimulation trials. Inside the parentheses are the values after removing delay outliers ($\tau_{sys} \geq 100\ ms$). The first row contains the $\tau_{sys}$ collected from all algorithms, followed by the delay values for individual algorithms. All the delays were measured in the unit of "sample." Since the sampling rate is 1 kHz, the resolution is 1 millisecond.

| | $\tau_{algo}$ | $\tau_{all}$ | $\tau_{sys}$ |
|---|---|---|---|
| overall | N/A | N/A | 32.11 ± 11.63 (31.95 ± 8.20) |
| AR | 4.57 ± 2.50 (4.57 ± 2.50) | 36.16 ± 8.17 (36.16 ± 8.17) | 31.60 ± 7.64 (31.60 ± 7.64) |
| ARFP | 7.46 ± 4.36 (7.45 ± 4.34) | 40.08 ± 15.63 (39.77 ± 9.98) | 32.62 ± 14.55 (32.31 ± 8.72) |

Measurement of $\tau_{algo}, \tau_{all}^{fake}, \tau_{sys}^{fake}$ in the Fake-Pulse System

The procedure for measuring $\tau_{algo}, \tau_{all}^{fake}$, and $\tau_{sys}^{fake}$ is as follows:

1. In the fake-pulse system, the processor continuously receives EEG data and predicts future phases upon the arrival of each new data chunk
2. For each prediction, record the computational time of the algorithm, $\tau_{algo}$, using the *tic* and *toc* functions in MATLAB.
3. When a target is detected, record the index of the current EEG sample (t=0) in the processor and send the marker.
4. $\tau_{all}^{fake}$ is calculated as the interval between the marker sending and receiving times in the processor.
5. $\tau_{sys}^{fake}$ is calculated as $\tau_{all}^{fake}$ minus $\tau_{algo}$.

The statistics below show the mean ± standard deviation of delay values over stimulation trials. Inside the parentheses are the values after removing delay outliers ($\tau_{sys}^{fake} \geq 50\ ms$). The first row contains the $\tau_{sys}^{fake}$ collected form all algorithms, followed by the delay values for individual algorithms. All the delays were measured in the unit of "sample." Since the sampling rate is 1 kHz, the resolution is 1 millisecond.

| | $\tau_{algo}$ | $\tau_{all}^{fake}$ | $\tau_{sys}^{fake}$ |
|---|---|---|---|
| overall | N/A | N/A | 26.24 ± 6.51 (26.19 ± 6.32) |
| AR | 4.51 ± 3.18 (4.51 ± 3.18 | 30.88 ± 7.19 (30.85 ± 7.10) | 26.37 ± 6.44 (26.34 ± 6.34) |
| ETP | 2.76 ± 0.34 (2.76 ± 0.34) | 29.05 ± 6.33 (29.02 ± 6.26) | 26.28 ± 6.32 (26.25 ± 6.24) |
| FFT | 2.83 ± 0.27 (2.83 ± 0.27) | 28.83 ± 6.35 (28.80 ± 6.31) | 26.00 ± 6.33 (25.98 ± 6.29) |
| PR | 2.83 ± 0.81 (2.83 ± 0.81) | 28.67 ± 6.69 (28.62 ± 6.53) | 25.84 ± 6.60 (25.80 ± 6.44) |
| ARFP | 6.90 ± 0.93 (6.89 ± 0.72) | 33.62 ± 7.09 (33.49 ± 6.30) | 26.71 ± 6.81 (26.60 ± 6.25) |

### S.2 Algorithm Parameter Optimization

Filter Selection

For fair performance comparison, all the algorithms should suffer the same amount of filter edge effect. Therefore, all the causal algorithms share the same FIR

bandpass filter which is also used to obtain the acausal ground truth.
The filter is designed by the MATLAB function

```
D=designfilt('bandpassfir', 'FilterOrder', 550, 'CutoffFrequency1',8,
'CutoffFrequency2', 13, 'SampleRate', 1000, 'DesignMethod', 'window');
```

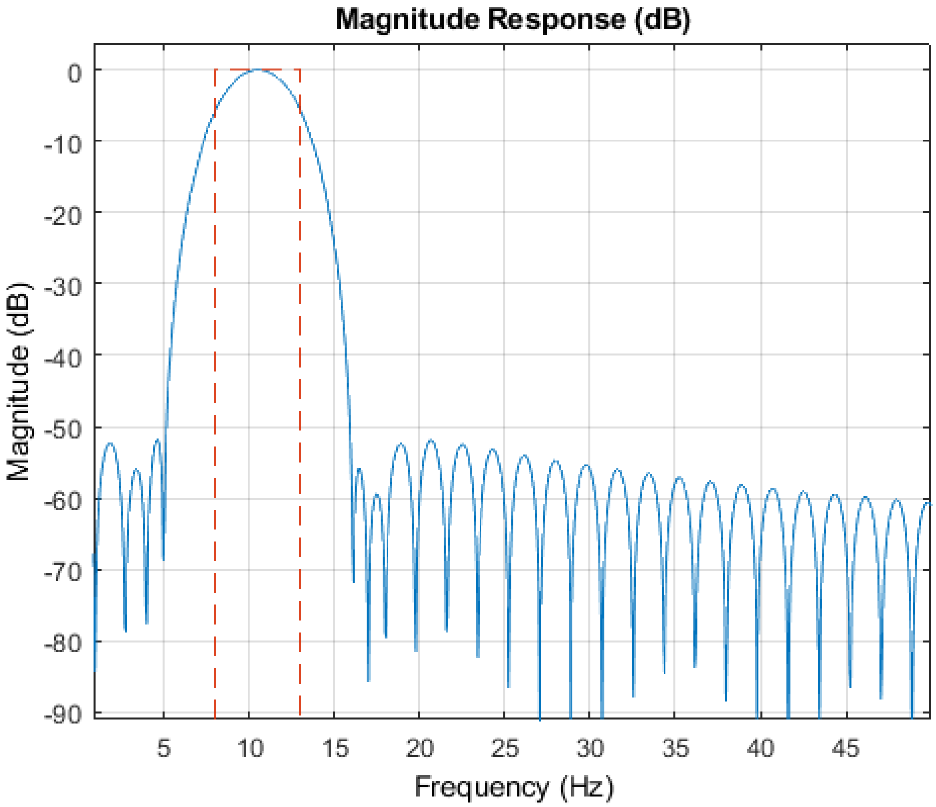


Note that the causal FIR filter used in the real-pulse validation was different from the above filter in order to reduce the group delay. The filter is designed by the MATLAB function.

```
D=designfilt('bandpassfir', 'FilterOrder', 140, 'CutoffFrequency1',8,
'CutoffFrequency2', 13, 'SampleRate', 1000);
```

Parameter Optimization

All the algorithm uses the parameters optimized by genetic algorithm (GA) except for ETP because the original paper offered the optimization method (two-stage calibration), and the GA-optimized parameters are not as good as the ones optimized by the original method for ETP.

1. AR

parameter list: $\mathbf{x} = [\text{window length} \quad \text{removed edge length} \quad \text{AR order}]^{\mathrm{T}}$

(window length: input signal is truncated to this length as the first step.

removed edge length: removed portion of bandpass-filtered signal at both ends before fitting the predictive model.)

fitness function: $\frac{1}{N}\sum_{n=1}^{N}\left|Arg\left(e^{j(\phi_{e,n}-\phi_{a,n})}\right)\right|$.

$N$: $number\ of\ training\ epochs.\ \phi_{e,n}$: $estimated\ current\ phase.\ \ \phi_{a,n}$: $ground\ truth$

constraint: $[-1 \quad 2 \quad 1]\mathbf{x} \leq -\mathbf{1}$

bound: $1651 \leq \mathrm{x}_1 \leq 2000, 1 \leq \mathrm{x}_2 \leq 120, 5 \leq \mathrm{x}_3 \leq 100$. ($\mathrm{x}_k$ denotes the $k^{th}$ element of $\mathbf{x}$.)

2. ETP

parameter list: $\mathbf{x} =$
$[\text{interpeak interval} \quad \text{minimum interpeak interval} \quad \text{window length} \quad \text{removed edge length}]^{\mathrm{T}}$

The parameters are optimized by the two-stage calibration proposed in the original paper.

3. FFT

parameter list: $\mathbf{x} = [\text{window length} \quad \text{removed edge length}]^{\mathrm{T}}$

fitness function: $\frac{1}{N}\sum_{t=1}^{T}\widetilde{w}(t)\sum_{n=1}^{N}\left|Arg\left(e^{j\left(\phi_{p,n}(t)-\phi_{a,n}(t)\right)}\right)\right|$.

$N$: $number\ of\ training\ epochs.\ T$: $number\ of\ future\ samples.$
$\phi_{p,n}(t)$: $predicted\ phase\ at\ t^{th}\ second.\ \ \phi_{a,n}(t)$: $ground\ truth\ at\ t^{th}\ second.$

$\widetilde{w}(t) = \frac{w(t)}{\sum_{t=1}^{T} w(t)}$, where $w(t) = \frac{1}{1+0.5\times(10^{-15})^{-\frac{t-0.005}{1000}}}$.

constraint: $[-1 \quad 2]\mathbf{x} \leq -\mathbf{1}$

bound: $1651 \leq \mathrm{x}_1 \leq 2000, 1 \leq \mathrm{x}_2 \leq 120$

4. PR

parameter list: $\mathbf{x} =$
$[\text{window length of signal} \quad \text{removed edge length of phase} \quad \text{window length of phase}]^{\mathrm{T}}$
.

(window length of signal: input signal is truncated to this length as the first step, after which the phases are calculated using the Hilbert-transform (HT)-based approach. removed edge length of phase: removed portion of phase data at both ends. window length of phase: the length of the remaining data to fit the linear regression model.)

fitness function: $\frac{1}{N}\sum_{n=1}^{N}\left|Arg\left(e^{j(\phi_{e,n}-\phi_{a,n})}\right)\right|$.

$N$: $number\ of\ training\ epochs.\ \phi_{e,n}$: $estimated\ current\ phase.\ \ \phi_{a,n}$: $ground\ truth$

constraint: $[-1 \quad 1 \quad 1]\mathbf{x} \leq \mathbf{0}$

bound: $1651 \leq \mathrm{x}_1 \leq 2000, 20 \leq \mathrm{x}_2 \leq 200, 5 \leq \mathrm{x}_3 \leq 360$

5. ARFP

parameter list: $\mathbf{x} =$
$[\text{window length} \quad \text{removed edge length} \quad \text{AR order} \quad \text{DC interval}]^{\mathrm{T}}$.

(DC interval: DC levels of padded signal and raw EEG within this interval are shifted to the same value before concatenation.)

fitness function: $\frac{1}{N}\sum_{n=1}^{N}\left|Arg\left(e^{j(\phi_{e,n}-\phi_{a,n})}\right)\right|$.

$N$: $number\ of\ training\ epochs.$ $\phi_{e,n}$: $estimated\ current\ phase.$ $\phi_{a,n}$: $ground\ truth$

constraint: $[-1\quad 2\quad 1\quad 0]\mathbf{x} \leq -\mathbf{1}$

bound: $1651 \leq \mathrm{x}_1 \leq 2000, 1 \leq \mathrm{x}_2 \leq 120, 5 \leq \mathrm{x}_3 \leq 100, 1 \leq \mathrm{x}_4 \leq 100$

### S.3 Distribution Fitting

Originally, $\tau_{all} = \tau_{sys} + \tau_{algo}$ and $\tau_{all}^{fake} = \tau_{sys}^{fake} + \tau_{algo}$. However, since the mean and variance of $\tau_{algo}$ are small compared to the intrinsic system delay ($\tau_{sys}$ and $\tau_{sys}^{fake}$), we assume the random variables $\tau_{all}$ and $\tau_{all}^{fake}$ to be the sum of the intrinsic system delay and the mean of $\tau_{algo}$. Specifically, the following two equations are assumed: $\tau_{all} = \tau_{sys} + \mathbb{E}(\tau_{algo})$ and $\tau_{all}^{fake} = \tau_{sys}^{fake} + \mathbb{E}(\tau_{algo})$, where $\mathbb{E}$ represents the expectation value. Therefore, $\tau_{all}$ and $\tau_{all}^{fake}$ can be obtained by first sampling from the fitted distributions of $\tau_{sys}$ and $\tau_{sys}^{fake}$, and then adding the expected value of $\tau_{algo}$. The following shows the fitting results of $\tau_{sys}$ and $\tau_{sys}^{fake}$.

$\tau_{sys}$

The distfit toolbox select the best fitting distribution as the one having the highest bootstrap score (*exponpow* in this case). However, based on the residual sum of squares (RSS), bootstrap score, and QQ plot, we consider the Weibull maximum (*weibull_max*) distribution (c=279.4450087362111, loc=1824.95969, scale=1796.681672) the best fitting one.

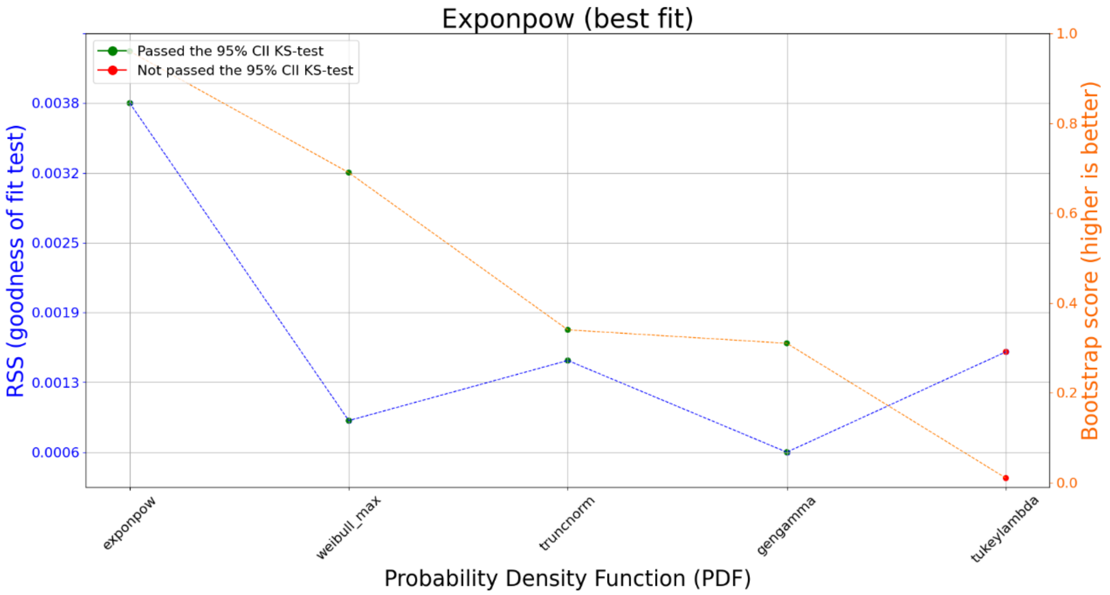
Exponpow (best fit)
Passed the 95% CII KS-test
Not passed the 95% CII KS-test
RSS (goodness of fit test)
Bootstrap score (higher is better)
0.0038
0.0032
0.0025
0.0019
0.0013
0.0006
1.0
0.8
0.6
0.4
0.2
0.0
exponpow
weibull_max
truncnorm
gengamma
tukeylambda
Probability Density Function (PDF)

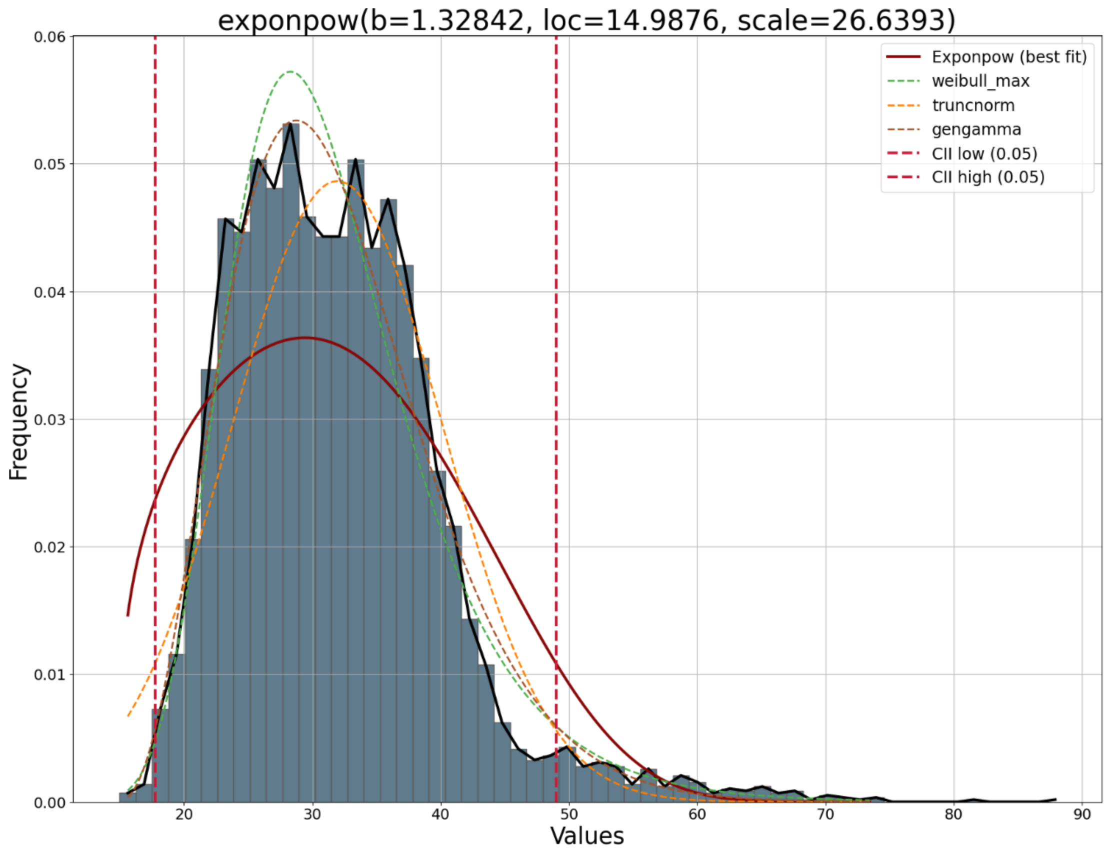
exponpow(b=1.32842, loc=14.9876, scale=26.6393)
Exponpow (best fit)
weibull_max
truncnorm
gengamma
CII low (0.05)
CII high (0.05)
Frequency
0.06
0.05
0.04
0.03
0.02
0.01
0.00
20
30
40
50
60
70
80
90
Values

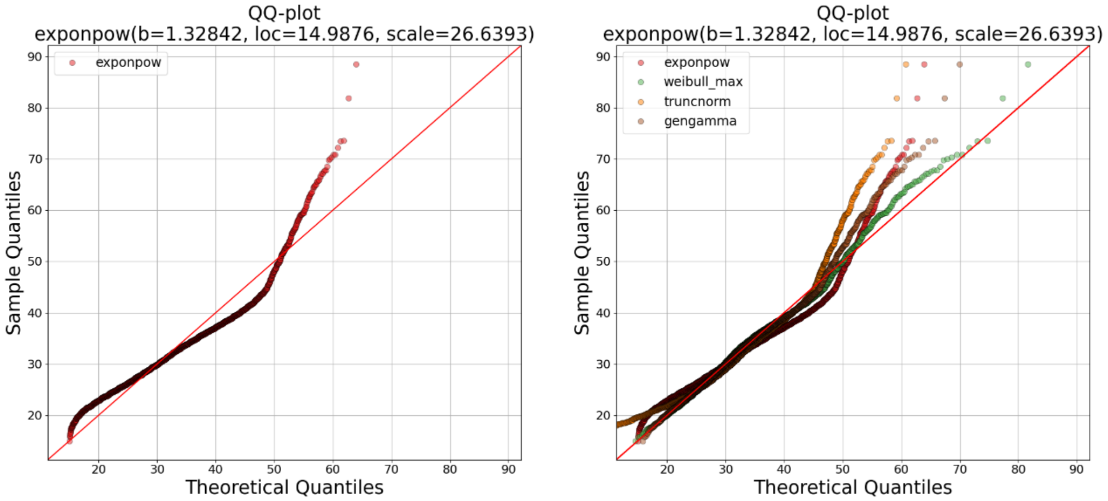


$\tau_{sys}^{fake}$

The distfit toolbox select the best fitting distribution as the one having the highest bootstrap score (*foldnorm* in this case). However, based on the residual sum of squares (RSS), bootstrap score, and QQ plot, we consider the power log-normal (*powerlognorm*) distribution (c=1.831511460745964, s=0.2116382697538689, loc=-9.471247, scale=39.032992) the best fitting one.

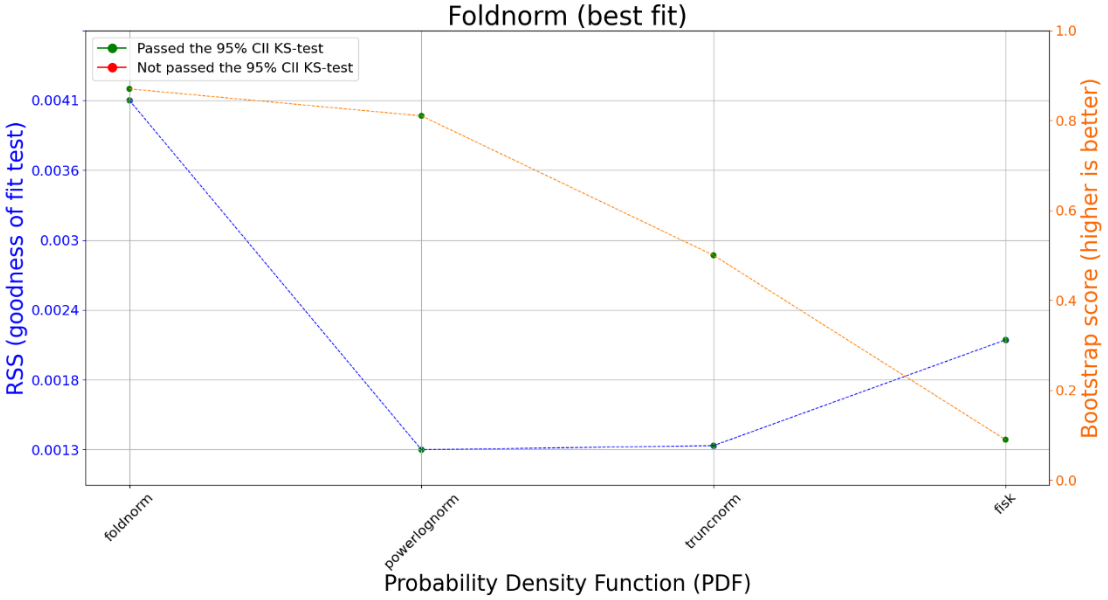

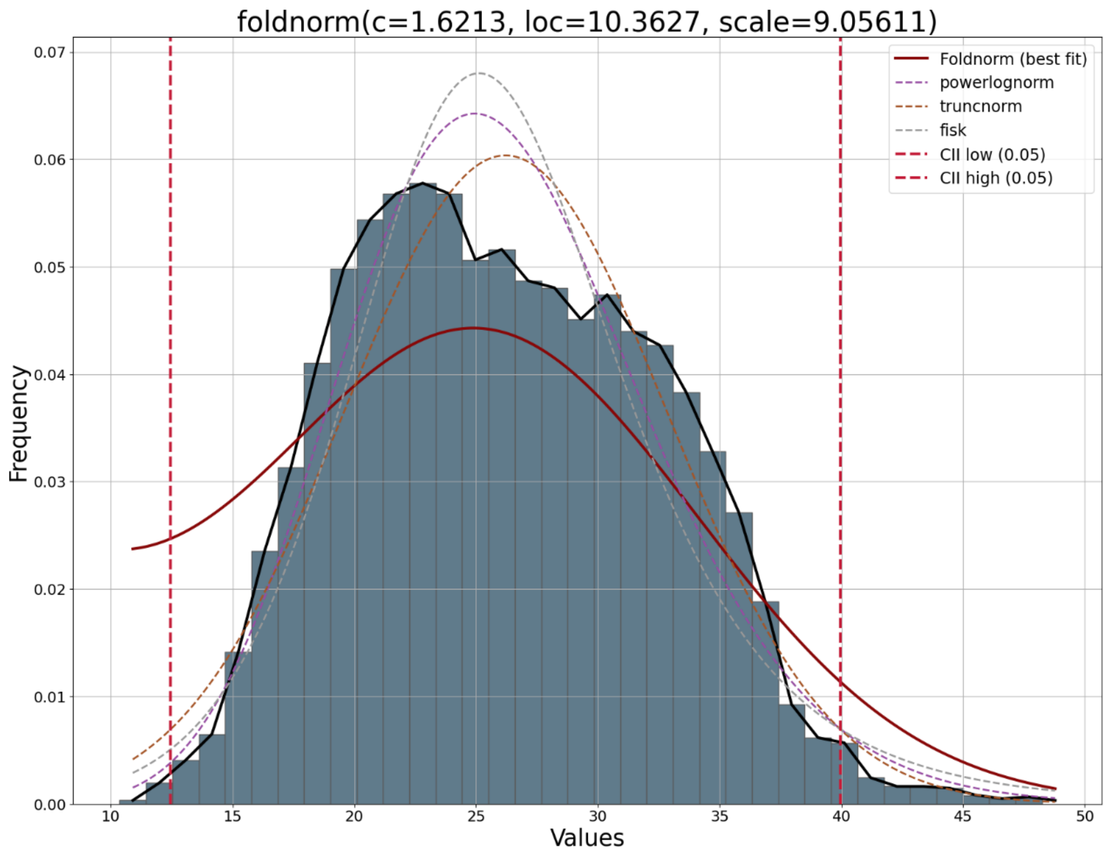
foldnorm(c=1.6213, loc=10.3627, scale=9.05611)
Frequency
Values
Foldnorm (best fit)
powerlognorm
truncnorm
fisk
CII low (0.05)
CII high (0.05)
0.00
0.01
0.02
0.03
0.04
0.05
0.06
0.07
10
15
20
25
30
35
40
45
50

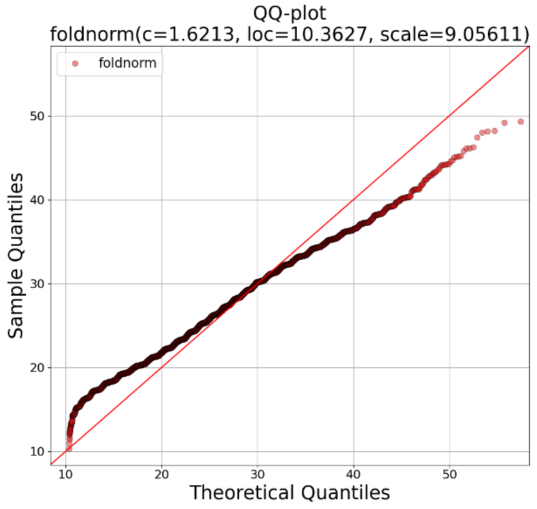
QQ-plot
foldnorm(c=1.6213, loc=10.3627, scale=9.05611)
foldnorm
Sample Quantiles
Theoretical Quantiles
10
20
30
40
50

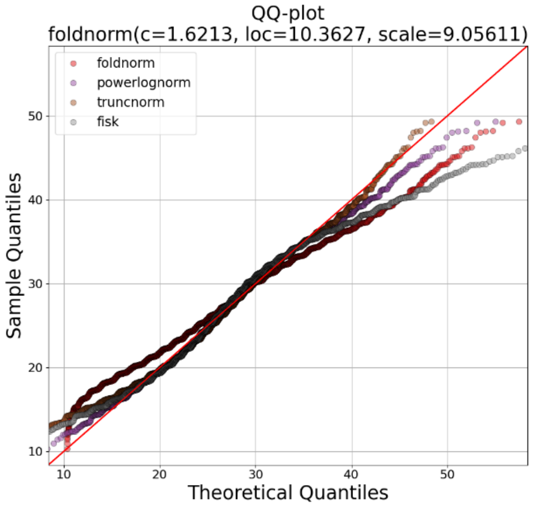
QQ-plot
foldnorm(c=1.6213, loc=10.3627, scale=9.05611)
foldnorm
powerlognorm
truncnorm
fisk
Sample Quantiles
Theoretical Quantiles
10
20
30
40
50

### S.4 Data Exclusion in Each Experiment

| Experiment | Data | Excluded sessions | Excluded trials |
|---|---|---|---|
| Predictive Performance and Robustness Evaluation | Fake-pulse | All sessions from s18 | None |
| In-silico validation | Fake-pulse | All sessions from s18 | Skipped trials / delay outliers |
| Fake-pulse validation | Fake-pulse | All sessions from s18 | Skipped trials (except for Fig. 5b in Main Text)/ delay outliers |
| Real-pulse validation | Real-pulse | All sessions from s18 | Skipped trials/ delay outliers |
| Equivalence of Validation Methods | Fake-pulse | All sessions from s18 | Skipped trials/ delay outliers |
| UBPP and Deadlock Rate | Fake-pulse | 1 session from s18 (data incompletion) | None |
| Distribution Fitting of $\tau_{sys}$ | Real-pulse | All sessions from s18 | None |
| Distribution Fitting of $\tau_{sys}^{fake}$ | Fake-pulse | All sessions from s18 | None |

Notes:

1. Skipped trials are the forcefully terminated stimulation trials because the target phase cannot be detected in 10 seconds.
2. Delay outliers are the stimulation trials with overlong $\tau_{sys}$ ($\geq 100\ ms$) or $\tau_{sys}^{fake}$ ($\geq 50\ ms$)

### S.5 Validation Framework for Timer Strategy

The following slides present the conceptual design of the validation framework for the timer-strategy.

Key Equation

- Validation framework for timer strategy

$$\tau_{timer}^{n} = \tau_{target}^{n} - \mathbb{E}(\tau_{all})$$

- Notations:
  $n$: trial index
  $\tau_{target}^{n}$: predicted target location
  $\tau_{pulse}^{n}$: actual pulse location
  $\tau_{marker}^{n}$: marker location
  $\phi_p$: predicted phase
  $\phi_a$: ground truth

Distribution Fitting

- To determine the distribution type of $\tau_{pulse}$, fix the $\tau_{timer}$ across $N$ stimulation trials in the real-pulse session to the same value and collect $\{\tau_{pulse}^{n}\}_{n=1,2,\ldots,N}$ for distribution fitting. The following paragraphs assume $\tau_{pulse}$ follows Weibull max distribution $WB(mean = \tau_{target}^{n}, variance = \sigma_{pulse}^{2})$.
- The distribution type of $\tau_{marker}$ can be determined using the same manner in the fake-pulse session. The following paragraphs assume $\tau_{marker}$ follows power log-normal distribution $PLN(mean = \tau_{timer}^{n} + \mathbb{E}(\tau_{all}^{fake}),\ variance = \sigma_{marker}^{2})$.

In-Silico Validation

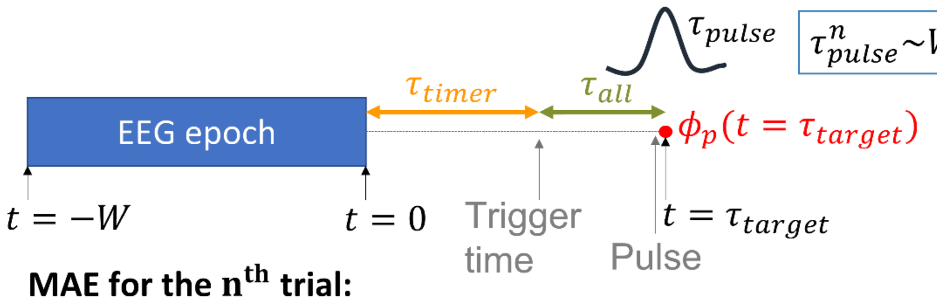


$\tau_{pulse}^{n} \sim WB(\tau_{target}^{n}, \sigma_{pulse}^{2})$

**MAE for the n^th trial:**

$$MAE_n = \sum_{m=1}^{M} \left|\arg\left(e^{j(\phi_p(t=\tau_{target}^{n}) - \phi_a(t=\tau_{pulse}^{n,m}))}\right)\right|),$$

$$where\ \tau_{pulse}^{n,m}\ are\ sampled\ from\ WB(\tau_{target}^{n}, \sigma_{pulse}^{2})$$

**MAE averaged across trials:**

$$MAE = \frac{1}{N}\sum_{n=1}^{N} MAE_n$$

Fake-Pulse Validation

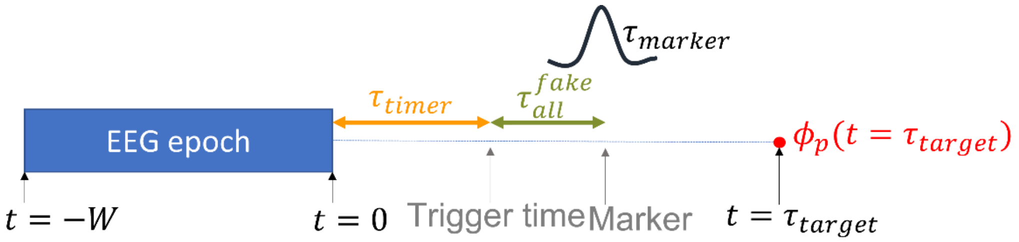


$\tau_{marker}^{n} \sim PLN\left(\tau_{timer}^{n} + \mathbb{E}\left(\tau_{all}^{fake}\right), \sigma_{marker}^{2}\right)$
$F$: quantile mapping from $PLN\left(\tau_{timer}^{n} + \mathbb{E}\left(\tau_{all}^{fake}\right), \sigma_{marker}^{2}\right)$ to $WB(\tau_{target}^{n}, \sigma_{pulse}^{2})$

**MAE for the n^th trial:**

$$MAE_n = \left|\arg\left(e^{j(\phi_p(t=\tau_{target}^{n}) - \phi_a(t=F(\tau_{marker}^{n})))}\right)\right|$$

**MAE averaged across trials:**

$$MAE = \frac{1}{N}\sum_{n=1}^{N} MAE_n$$

Real-Pulse Validation

The procedure is the same as the trigger-range strategy (average waveform prior to pulse onset)

### S.6 Equivalence of Validation Methods

| Validation Methods | System | Stimulus Type | EEG data | Groud Truth | Performance Metric |
|---|---|---|---|---|---|
| in-silico | computer simulation | simulated pulse locations | prerecorded resting-state EEG | available | quantitative |
| fake pulse | fake-pulse system | event markers | online EEG with event markers | available | quantitative |
| real pulse | real pulse system | TMS pulses | online EEG with TMS pulses | unavailable | qualitative |

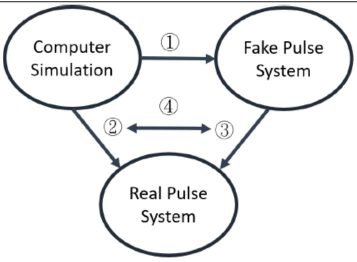


① In-silico validation to estimate fake-pulse accuracy (justification of in-silico validation)
② In-silico validation to estimate real-pulse accuracy
③ Fake pulse validation to estimate real-pulse accuracy
④ Compare the MAE of ② and ③ (justification of fake pulse validation)

| | Target | AR | ETP | FFT | PR | ARFP |
|---|---|---|---|---|---|---|
| Step1 | non-phase-locked | 0.4939 | 0.4065 | 0.7922 | 0.4896 | 0.7245 |
| Step4 | 0° | 0.3233 | 0.1061 | 0.4133 | 0.5174 | 0.2543 |
| | 90° | 0.1653 | 0.3199 | 0.7173 | 0.6357 | 0.0087** |
| | 180° | 0.6108 | 0.0867 | 0.7841 | 0.7218 | 0.6443 |
| | 270° | 0.2789 | 0.1870 | 0.4024 | 0.9917 | 0.2281 |
| | pooled MAE | 0.3163 | 0.6691 | 0.6228 | 0.4450 | 0.1145 |

Figure S6: Equivalence of validation methods. The upper table summarizes the three validation methods that comprise the delay-relevant validation framework. The lower-left figure shows the procedures to justify the equivalence of the three methods. The lower-right table reports the p-value of step 1 and 4 in the lower-left figure. Except the 90° of ARFP, all the methods produce similar MAE. The results suggest that fake-pulse and in-silico validation are valid and equivalent surrogates for real-pulse validation (actual stimulation accuracy)

##### Methods

As TMS artifacts disabled the calculation of ground truth, either in-silico or fake-pulse validation was utilized to represent the accuracy of actual TMS outcomes. To affirm the equivalence of in-silico and fake-pulse validation as valid surrogates for real-pulse validation, the following steps were carried out (see Figure S6 above):

Step 1. Justification of in-silico validation.
Step 2. Estimation of real-pulse accuracy through in-silico validation.
Step 3. Estimation of real-pulse accuracy through fake-pulse validation.
Step 4. Justification of fake-pulse validation.

To ensure that the difference in MAE was solely attributable to methodological variations rather than data discrepancies, the epoched data employed in the in-silico approach precisely matched the input data provided to the algorithm during the real-time execution in the fake-pulse session.

In Step 1, in order to substantiate the capability of in-silico validation in emulating real-time scenarios, Wilcoxon signed-rank test was applied to compare the $MAE_n$ derived from the in-silico approach (in-silico MAE) with the $MAE_n$ calculated from the actual marker locations (marker-based MAE) in fake-pulse sessions. To compute the in-silico MAE, marker locations, denoted as $\tau_{all}^{fake}$, were simulated by adding $\mathbb{E}(\tau_{algo})$ to randomly sampled points from the best-fitting power-log-normal distribution as described in Section 2.4.2 of the Main Text. After that, Equation 3 in the Main Text was modified to the form of $MAE_n(\mathbb{E}(\tau_{all}^{fake}), \mathbb{V}(\tau_{all}^{fake}))$ to compute the error between the predicted phase and the ground truth at the simulated marker locations. Here, $\mathbb{E}(\tau_{all}^{fake})$ and $\mathbb{V}(\tau_{all}^{fake})$ represent the sample mean and sample variance of $\tau_{all}^{fake}$, respectively. In contrast, the marker-based MAE for each trial was calculated as follows:

$$MAE_n = \left|\arg(e^{i(\phi_{p,n}-\phi_a)})\right|$$

In the above equations, $\phi_{p,n}$ signifies the predicted phase for trial $n$ at $\mathrm{t} = \mathbb{E}(\tau_{all}^{fake})$, while $\phi_a$ represents the ground truth at the actual marker location ($\mathrm{t} = \tau_{all,n}^{fake}$). The Wilcoxon signed-rank test was applied to compare in-silico MAE and marker-based MAE. Because the triggering of markers during the fake-pulse sessions depended on the predicted phase at $\mathrm{t} = \mathbb{E}(\tau_{all})$ rather than at $\mathrm{t} = \mathbb{E}(\tau_{all}^{fake})$, the MAEs in Step 1 were not associated with any specific target phase.

In Step 2, in-silico validation (as defined in Equation 3 in the Main Text) was used to model the accuracy of the actual TMS pulses for each epoch. Likewise, in Step 3, fake-pulse validation (as defined in Equation 6 in the Main Text) was employed to estimate accuracy based on the adjusted marker location. In Step 4, a Wilcoxon signed-rank test was conducted to compare the MAE derived in Step 2 and Step 3.

#### Results

Figure S6 above shows no significant difference between in-silico and marker-based MAE in step 1. The results in step 4 reveal no significant difference between in-silico validation and fake-pulse validation, except for ARFP at 90°. To ensure that the mismatch in ARFP at 90° is not due to methodological error, we conducted a Kolmogorov-Smirnov test and confirmed that the adjusted marker locations did not deviate from the distribution of the actual pulse locations. The results in the following paragraph suggest that the mismatch in MAEs is a coincidence, where the fake-pulse MAE deviated in the same direction as the in-silico MAE in the low-MAE region. In conclusion, the results in step 1 and step 4 demonstrate that in-silico and fake-pulse validation are valid and equivalent surrogates to quantify the stimulation accuracy of real-pulse system.

#### Mismatch in ARFP at 90°

| | Target | AR | ETP | FFT | PR | ARFP |
|---|---|---|---|---|---|---|
| Step1 | non-phase-locked | 0.4939 | 0.4065 | 0.7922 | 0.4896 | 0.7245 |
| Step4 | 0° | 0.3233 | 0.1061 | 0.4133 | 0.5174 | 0.2543 |
| | 90° | 0.1653 | 0.3199 | 0.7173 | 0.6357 | **0.0087**** |
| | 180° | 0.6108 | 0.0867 | 0.7841 | 0.7218 | 0.6443 |
| | 270° | 0.2789 | 0.1870 | 0.4024 | 0.9917 | 0.2281 |
| | pooled MAE | 0.3163 | 0.6691 | 0.6228 | 0.4450 | 0.1145 |

For ARFP at 90°, in-silico and fake-pulse validation produced significantly different MAEs. We conducted a Kolmogorov-Smirnov test to investigate whether this difference results from a systematic error where the distributions of the actual data deviate from the best-fitting distributions. The results are shown below.

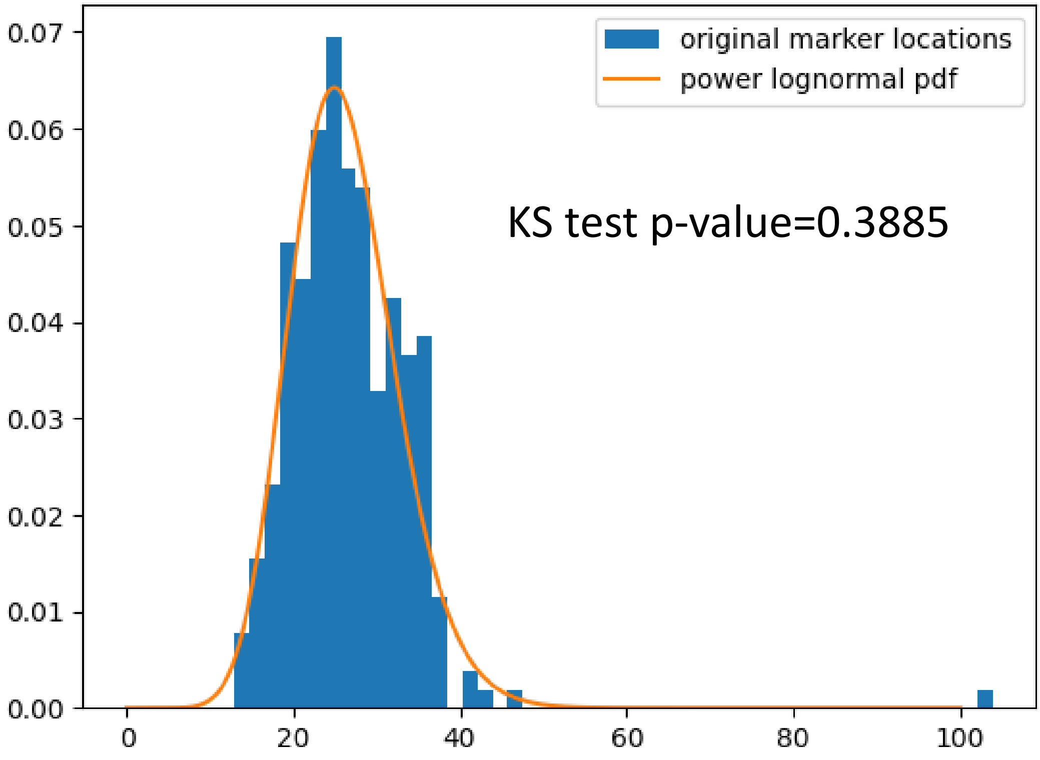


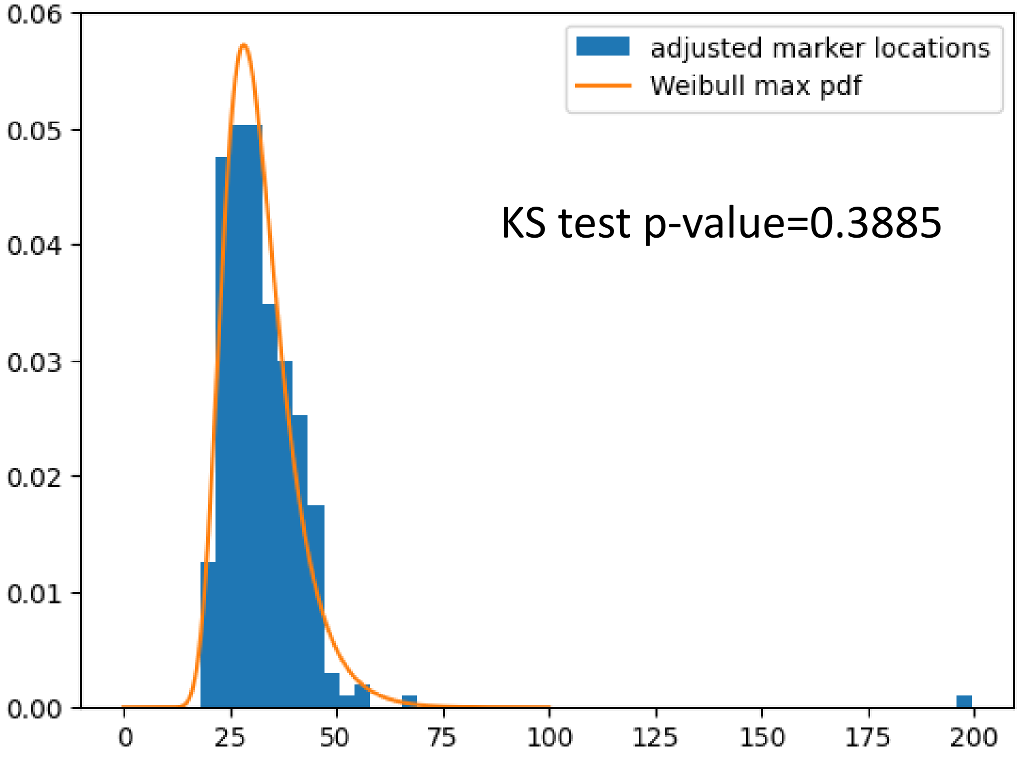

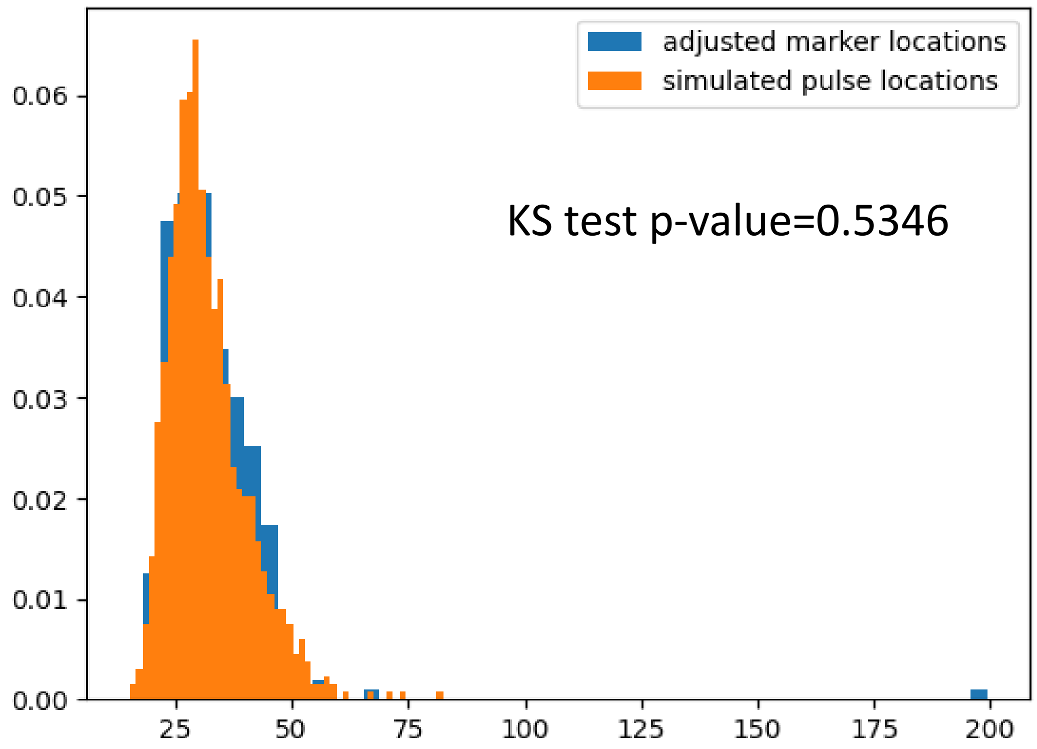


The results indicate that the simulated pulse locations in in-silico validation and the adjusted marker locations in fake-pulse validation do not significantly differ from each other. It is important to note that $MAE_n$ (Mean Absolute Error for each epoch) is computed from aggregate simulated pulse locations ($\tau_{all,1}$ to $\tau_{all,M}$) in in-silico validation, whereas it is derived from only one adjusted marker location in fake-pulse validation, possibly increasing the sampling error. Therefore, we hypothesize that the observed mismatch for ARFP at 90° is merely coincidental.

To investigate this further, the $MAE_n$ values from both validation methods are presented in the figure below. Each point in the figure corresponds to an epoch. The results demonstrate that the MAEs from both methods are generally comparable, except for some trials exhibiting notably low fake-pulse MAE. We attribute these instances of low fake-pulse MAE to incidental events, where the adjusted marker locations happen to fall at $\mathrm{t} = \mathbb{E}(\tau_{all})$. In contrast, in-silico validation simulates numerous pulses per trial, making such incidental events less likely to occur.

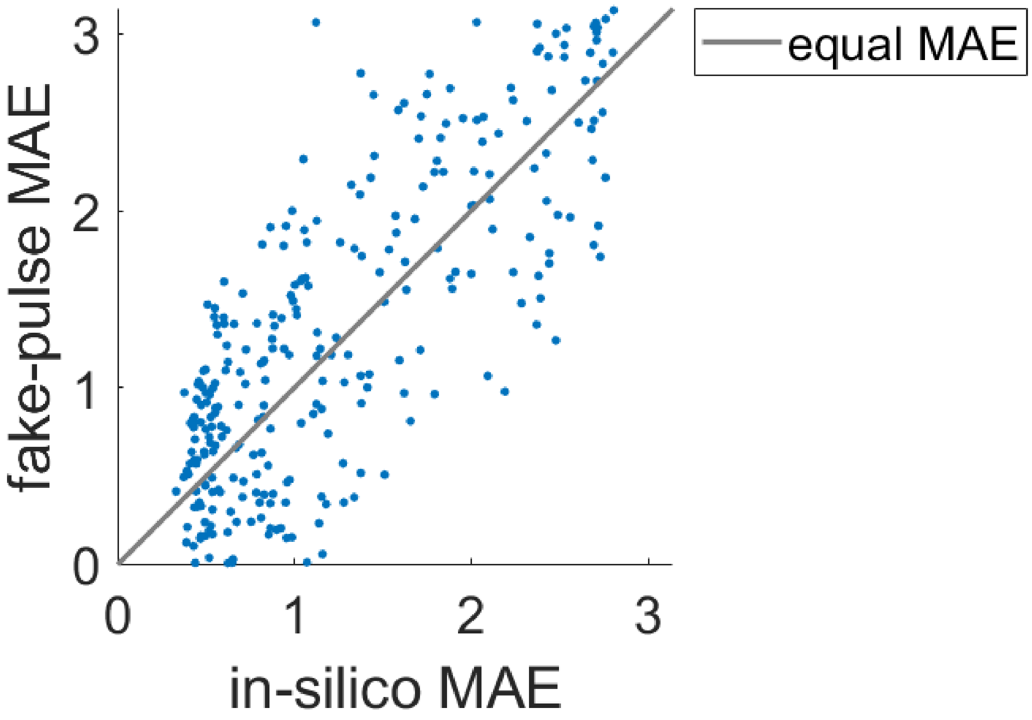


### S.7 UBPP and Deadlock

Methods

- IBF and deadlock rate for each algorithm

The IBF and deadlock rate of each algorithm are reported to evaluate their robustness. To calculate the IBF, 100 prediction attempts were randomly selected for each algorithm in each fake-pulse session. A polar histogram is used to visualize the predictive distribution at $t = 0$ and $t = \mathbb{E}(\tau_{all})$, along with the corresponding ground truth. The IBF, calculated from Equation 2 in the Main Text, is presented above each polar histogram. In contrast, the deadlock rate was computed by dividing the number of skipped trials (due to communication deadlock) of an algorithm by the number of stimuli associated with that algorithm in a session.

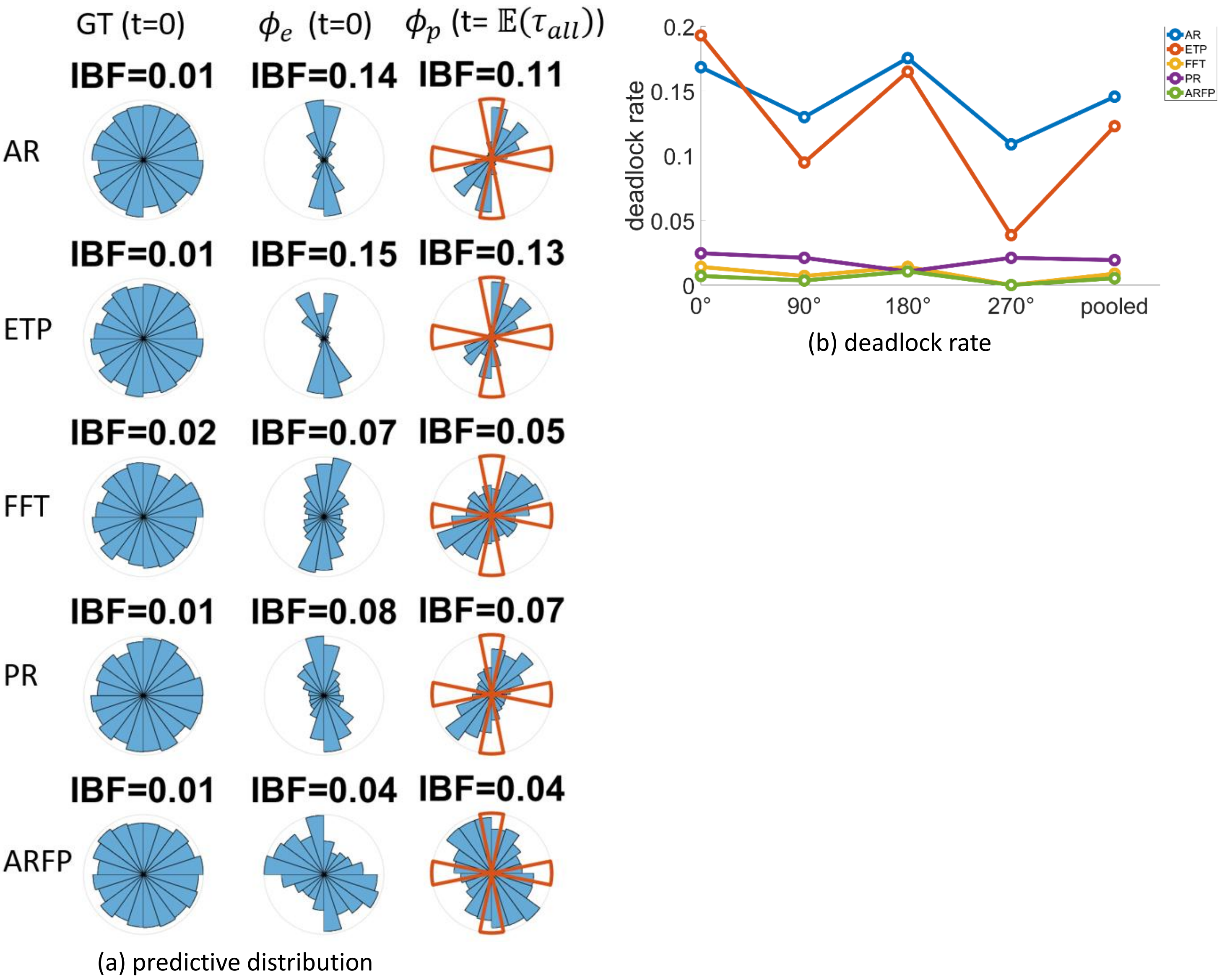


Figure S7: UBPP and deadlock. (a) Imbalance factor of different algorithms. The figure presents the ground truth at $t = 0$, estimated phase at $t = 0$, and the predicted phase at $t = \mathbb{E}(\tau_{all})$ in the left, middle, and right columns, respectively. The orange sectors in the right column represent the trigger range for the four target phases. (b) Deadlock rate or each of the four targets and the aggregate deadlock rate for all four targets.

- IBF and deadlock rate for each algorithm

Figure S7a depicts the predictive distributions of different algorithms. The results suggest that the UBPP introduced by zero padding at $\mathrm{t} = 0$ can persist throughout the prediction period until $\mathrm{t} = \mathbb{E}(\tau_{all})$, ultimately leading to communication deadlock. ARFP, with the aid of forecast padding, exhibits the lowest IBF and, consequently, the lowest deadlock rate among all algorithms (Figure S7b).

### S.8 Baseline Correction

Given the ability of forecast padding (FP) to reduce the edge effect of filtering and enhance the accuracy and robustness of phase prediction, we explored whether baseline correction before filtering could bring further enhancement. At first glance, baseline correction may appear redundant since bandpass filtering already rejects DC component. However, the default filtering process (*filter* and *filtfilt* function) in MATLAB automatically applies zero-padding before convolution, and causes a sharp transition at the end. Even for the bidirectional filtering (*filtfilt* function), where initial conditions are optimized to mitigate transient effect [20], the vanished amplitudes at both ends of signal are still significant. Therefore, we investigate the effects of the following three baseline correction (BC) methods on phase prediction:

1. No baseline correction (None): the EEG epoch is directly provided to the phase prediction algorithm without any baseline correction.
2. Standard baseline window (SBW): the mean of the entire epoch is subtracted from that epoch. In other words, the baseline window is equal to the length of the entire epoch.
3. Optimized baseline window (OBW): In the training stage, the baseline window length is optimized[1] for each algorithm. In the inference stage, the DC level is calculated by averaging the signal within the optimized baseline window around t=0. The DC level is then subtracted from the entire epoch before filtering.

1. In the training stage of QA phase estimation, grid search is applied to find the optimal baseline window length between 1 and 100 ms. In the training stage of AR, FFT, PR, and their forecast padding counterparts, the baseline window length and other parameters are optimized together using a genetic algorithm. For ETP and its forecast padding version, since the parameters are trained on continuous EEG, baseline correction plays no role in the training stage. Therefore, the baseline window used for the inference stage is chosen as 50 ms based on empirical rules.

To assess the efficacy of different BC methods, we first applied them to a newly proposed baseline method: quasi-acausal (QA) phase estimation. In this approach, the raw EEG epochs are zero-padded, and zero-phase bandpass filtering along with the Hilbert transform are applied to the padded signals to calculate the phase at $t = 0$. Since this method does not involve any predictive model, it represents the minimal processing steps in a phase prediction algorithm. The term "quasi-acausal" indicates that the processing steps are identical to those of the acausal phase estimation described in Preliminary Section 2, except that the future signal is replaced with padded zeros. The accuracy and robustness of the QA approach under different BC methods are presented below. The Kruskal-Wallis test followed by the Tukey-Kramer post hoc test demonstrates a significant effect of BC type on MAE and IBF of the QA approach

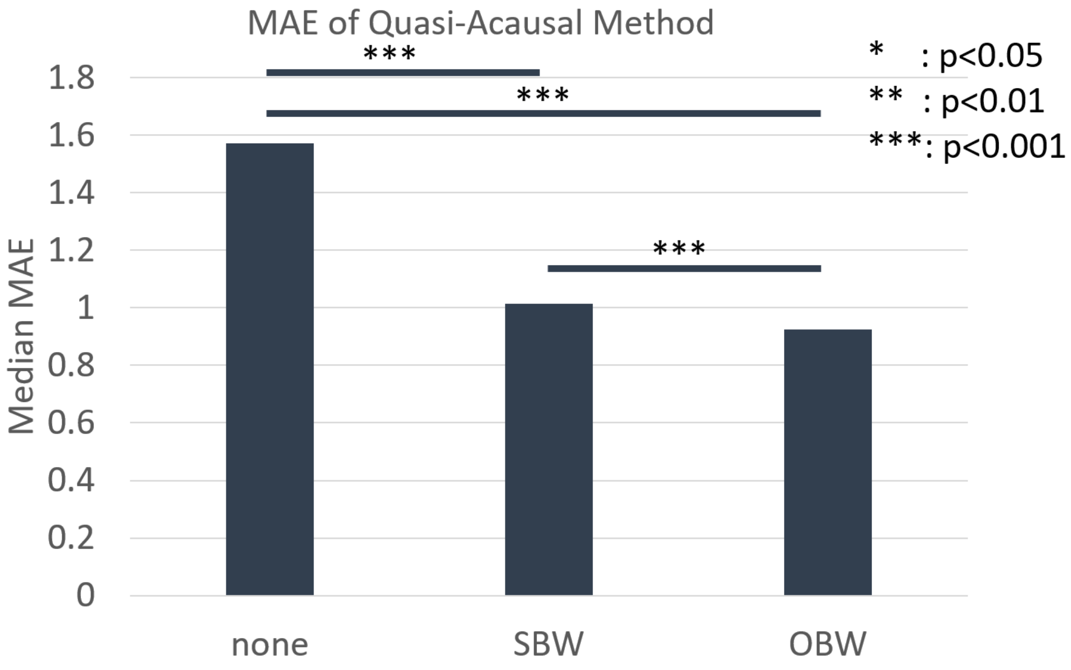

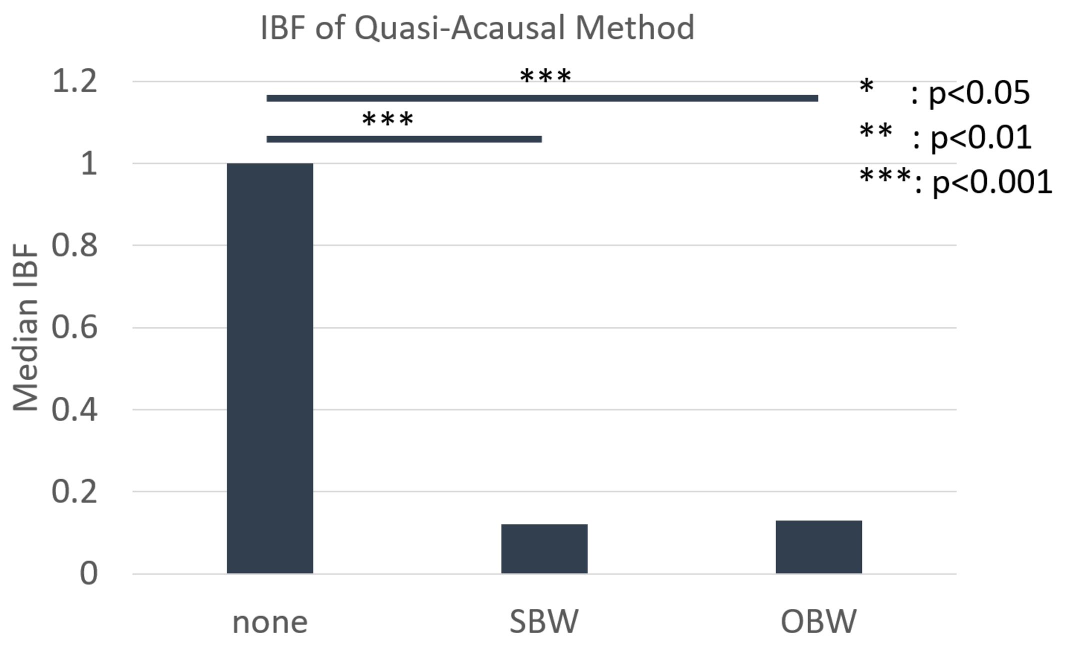


Next, we investigated the accuracy and robustness of all combinations of algorithms, forecast padding, and BC methods. Each BC method was simultaneously applied to both training and testing data to achieve optimal performance. The median MAE and IBF are presented below. A three-way ANOVA[2] shows a significant main effect of algorithm type and the application of forecast padding, while baseline correction has an insignificant effect.

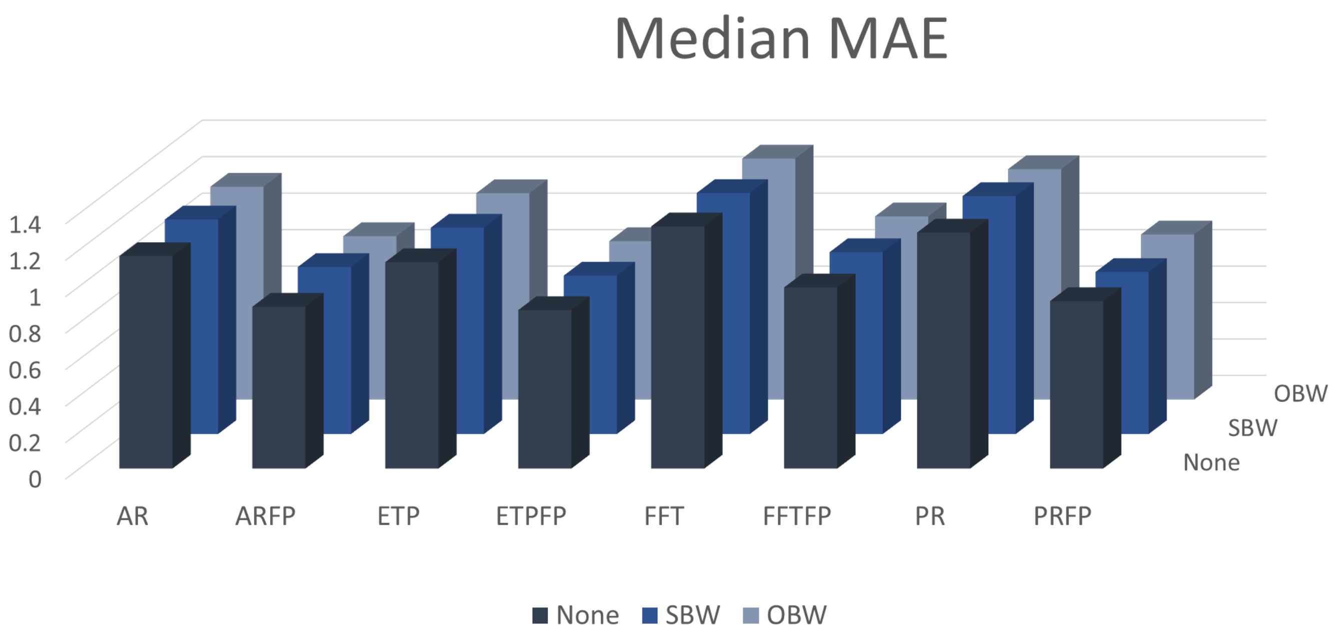


2. Before running the ANOVA, Box Cox transformation was employed to increase the Gaussianity of the MAE and IBF data. The three independent variables are BC method, algorithm type, and application of FP. The BC method has 3 levels (None, SBW, and OBW); the algorithm type has four levels (AR, ETP, FFT, and PR); the application of FP has two levels (true or false).

### ANOVA For MAE

| Source | Sum Sq. | d.f. | Mean Sq. | F | Prob>F |
|---|---|---|---|---|---|
| BC | 1 | 2 | 0.49 | 0.63 | 0.532 |
| Algo | 222.9 | 3 | 74.3 | 96.51 | 0 |
| FP | 1653 | 1 | 1652.98 | 2147.27 | 0 |
| BC:Algo | 1.9 | 6 | 0.31 | 0.41 | 0.8742 |
| BC:FP | 0.4 | 2 | 0.19 | 0.25 | 0.7775 |
| Algo:FP | 56.4 | 3 | 18.79 | 24.41 | 0 |
| BC:Algo:FP | 1.1 | 6 | 0.18 | 0.23 | 0.9666 |
| Error | 105290.5 | 136776 | 0.77 | | |
| Total | 107227 | 136799 | | | |

Constrained (Type III) sums of squares.

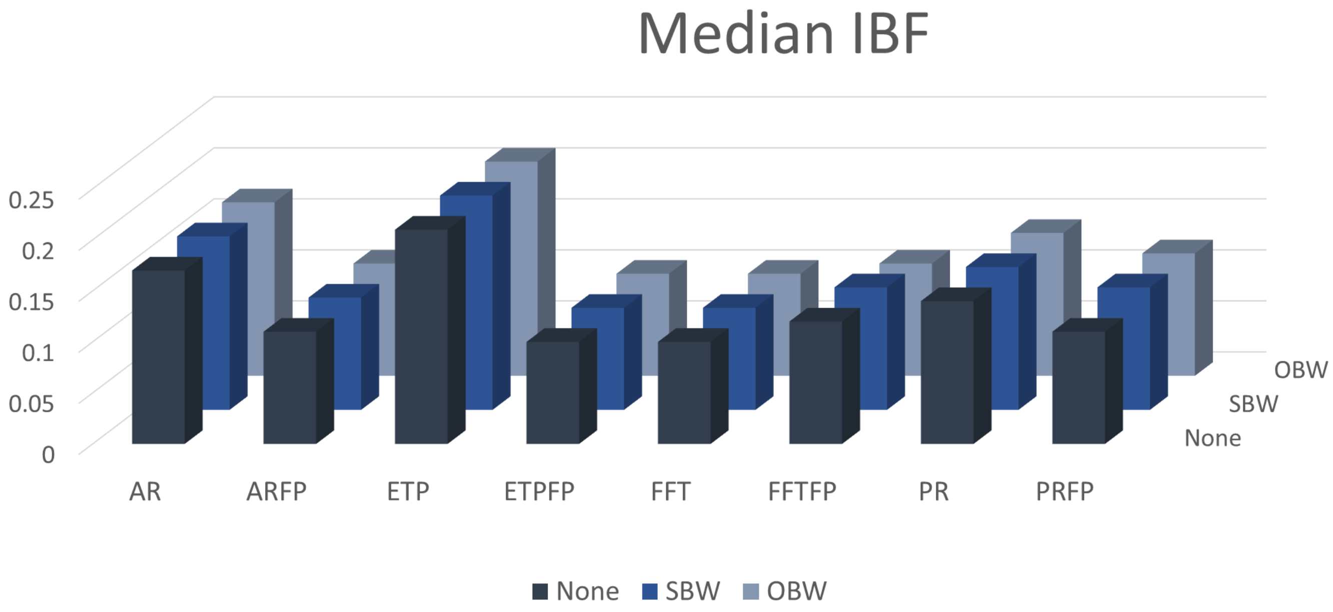

#### ANOVA For IBF

| Source | Sum Sq. | d.f. | Mean Sq. | F | Prob>F |
|---|---|---|---|---|---|
| BC | 2.88 | 2 | 1.44 | 0.75 | 0.4702 |
| Algo | 164.32 | 3 | 54.773 | 28.72 | 0 |
| FP | 202.76 | 1 | 202.763 | 106.33 | 0 |
| BC:Algo | 2.83 | 6 | 0.471 | 0.25 | 0.9606 |
| BC:FP | 1.24 | 2 | 0.622 | 0.33 | 0.7219 |
| Algo:FP | 227.01 | 3 | 75.669 | 39.68 | 0 |
| BC:Algo:FP | 1.91 | 6 | 0.319 | 0.17 | 0.9855 |
| Error | 2562.94 | 1344 | 1.907 | | |
| Total | 3165.89 | 1367 | | | |

Constrained (Type III) sums of squares.

### S.9 Prediction Gain

To investigate the utility of the predicted information, we compare the accuracy of the causal phase prediction algorithms with that of the QA method. The Prediction Gain (PG) is defined by the following equation:

$$PG_{MAE} = median\ MAE_{QA} - median\ MAE_{pred}$$

Here, $MAE_{QA}$ represents the MAE for the quasi-acausal method across all EEG epochs, and $MAE_{pred}$ represents the MAE for phase prediction algorithms across all EEG epochs. A PG value of 0 indicates that the predicted information adds no value beyond simply padding the signal with zeros. Positive PG values signify that the predictions enhance accuracy compared to zero-padding, while negative values suggest the predictions are detrimental. Similarly, the prediction gain for IBF is defined as: $PG_{IBF} = median\ IBF_{QA} - median\ IBF_{pred}$. The figures below illustrate the PG values for MAE and IBF across various combinations of algorithms, padding schemes, and BC strategies. Red bins represent negative PG, indicating less informative predictions than zero-padding, while blue bins show zero or positive PG values. Notably, for non-forecast padded algorithms, 8 out of 12 combinations exhibit negative PG for MAE and 6 out of 12 for IBF. In contrast, using forecast padding, where predicted signal complements the raw EEG, only 1 combination has negative PG for MAE and none for IBF.

These findings challenge the traditional practice in phase prediction of discarding distorted data and solely relying on predicted information. Our results demonstrate that forecast padding effectively harnesses the complementary nature of predicted data, maximizing its efficacy and ultimately leading to improved accuracy and robustness compared to zero-padding.

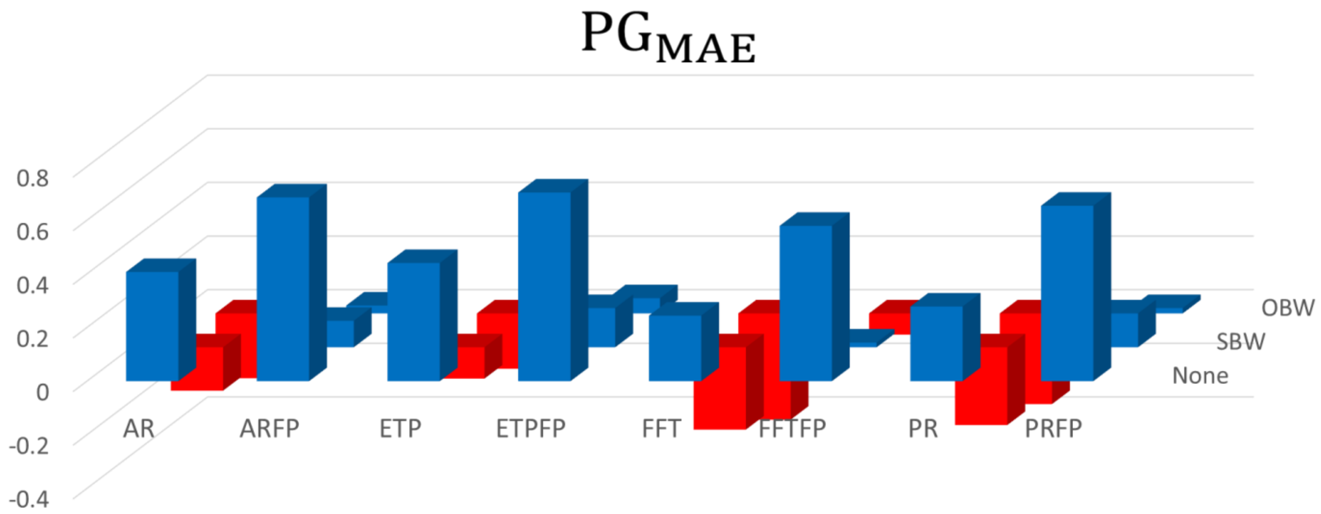


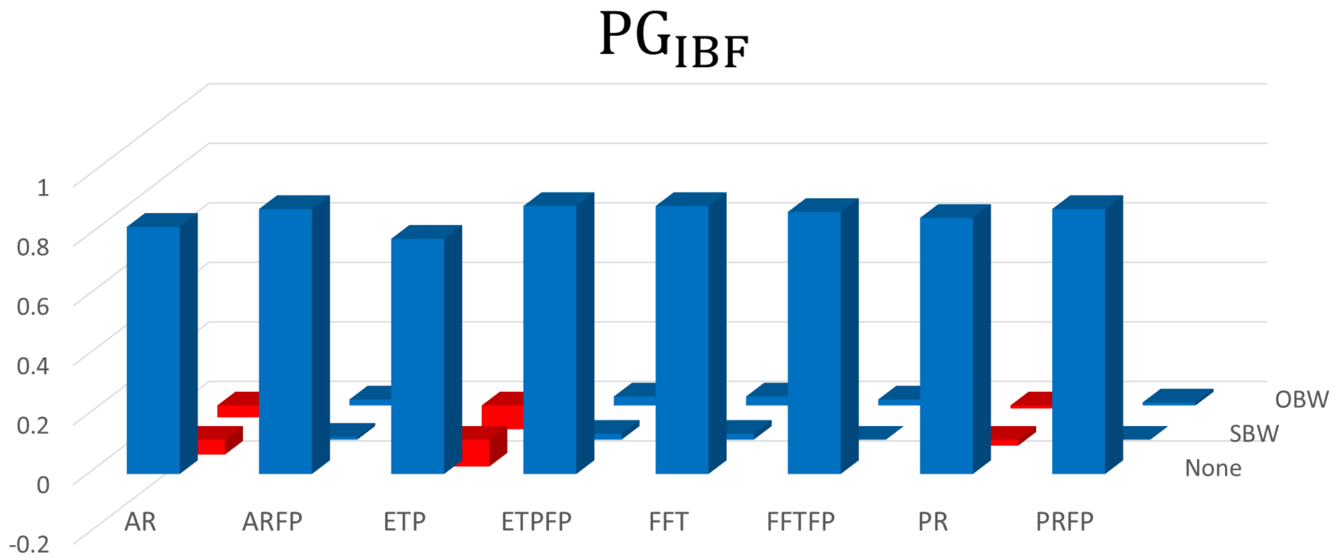

### S.10 Algorithm Implementation

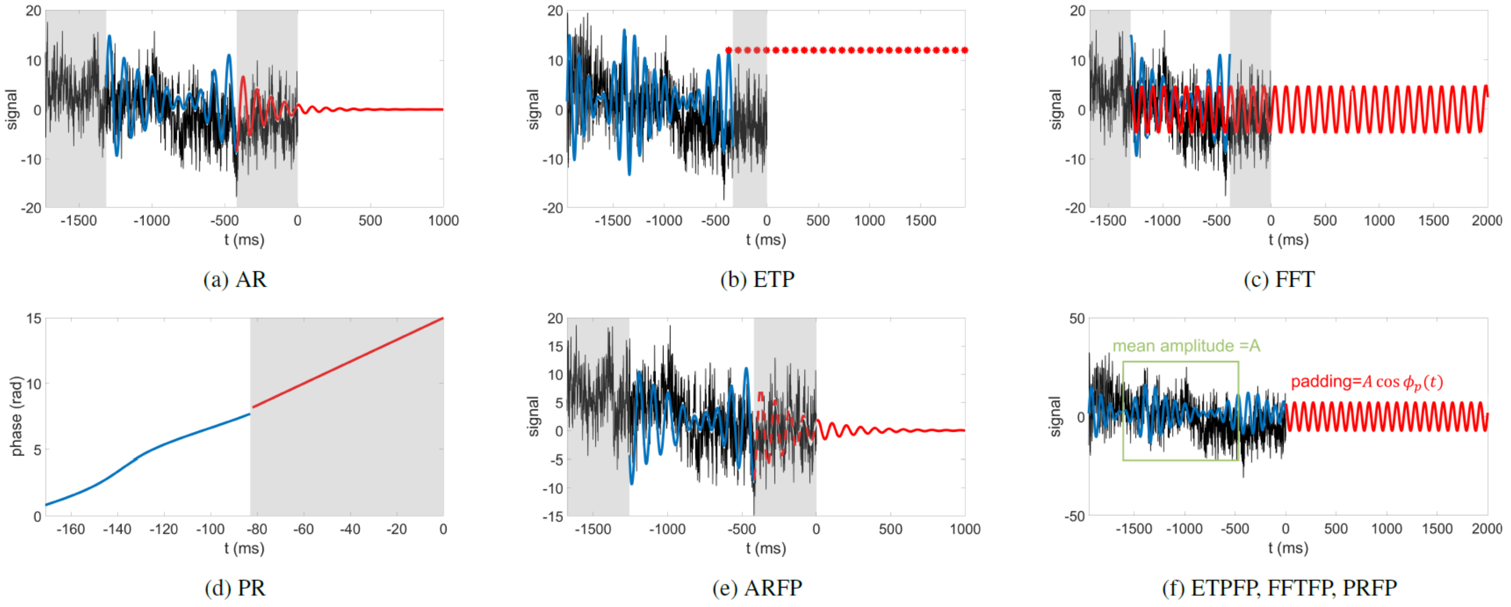


Figure S10: The diagram of three commonly used and one newly proposed phase prediction algorithms and their forecast-padded version.

We implemented four algorithms and their forecast padding version for performance evaluation, where AR [2], ETP [11] and FFT [12] are according to their original papers, and phase regression (PR) are our proposed methods. Five out of eight algorithms (AR, ETP, FFT, PR, and AR with forecast padding) are further implemented on the closed-loop TMS system to evaluate the stimulaton accuracy and deadlock rate. According to the design principle in our previous work [4], ETP and FFT are *pure prediction*, where the phase is directly predicted at $t = \tau_{all}$, and AR, PR, and all the forecast padded versions are *predictive estimation*, where the current phase at $t = 0$ is first estimated and the delay-relevant phase offset is added to predict the future phase. The phase offset can be obtained from the calibration data as follows.

$$\Delta\theta = \arg(\sum_n e^{i(\phi_{a,n}-\phi_{e,n})}) \quad \text{(S10-1)}$$

Here, $\phi_{a,n}$ is the acausally calculated ground truth of the $n^{th}$ calibration trial at $t = \tau_{all}$, $\phi_{e,n}$ is the estimated current phase at $t = 0$, and $\arg$ is the wrapped phase of the complex number between $-\pi$ and $\pi$. The parameters of AR, FFT, PR, and ARFP are optimized using the genetic algorithm (see Supplementary S.2) and the parameters of ETP are optimized using the two-stage calibration as in the original paper [11].

Autoregressive Phase Estimation (AR)

The overall process is illustrated in Figure S10a. The signal is zero-phase bandpass filtered within 8-13 Hz, then the edges at both ends (shaded gray regions) are removed due to filter edge effect. The remaining part (blue line) is used to fit the autoregressive (AR) model. The fitted AR coefficients are then used to iteratively predict future samples by linearly combining the previous samples. Next, the forecast signal (red line) is concatenated to the bandpass filtered signal (including the beginning edge portion). Finally, the concatenated signal is Hilbert transformed to estimate the current phase at $t = 0$. Phase offset (Equation S10-1) is added to the estimated current phase to predict future phases.

Educated Temporal Prediction (ETP)

The overall process is illustrated in Figure S10b. The signal is zero-phase bandpass filtered within 8-13 Hz, then the edge (shaded gray region) is removed due to filter edge effect. The peaks are identified as the local maxima in the remaining signal (blue line). Multiples of the inter-peak interval obtained in the calibration stage are added to the latest peak to forecast the future peaks (red asterisks). Phases between two adjacent peaks are linearly interpolated in the form of

$\phi(t) = \frac{2\pi t}{T}$, where $T$ is the inter-peak interval and $t$ is the time since the last peak.

Fast Fourier Transform (FFT)

The overall process is illustrated in Figure S10c. The signal is zero-phase bandpass filtered within 8-13 Hz, then the edges at both ends (shaded gray regions) are removed due to filter edge effect. The FFT of the remaining part (blue line) is calculated and the frequency bin of the largest magnitude within the alpha band is chosen as the dominant frequency. The dominant frequency and its corresponding phase (angle of the complex FFT signal) are used to construct the cosine signal (red line). Finally, the phase of the cosine signal at $t = \tau_{all}$ is extrapolated.

Phase Regression (PR)

This algorithm is inspired by the fact that the phase of a sine wave is a linear function of time. For the narrowband signal, the phase-time relationship is approximated using linear least square regression. The overall process is illustrated in Figure S10d. The signal is zero-phase bandpass filtered within 8-13 Hz, and the Hilbert transform is applied to calculate the instantaneous phase. The edge portion (shaded gray region) is removed due to filter edge effect. A window of the remaining portion (blue line) is used to fit the linear model using the least-square method. The linear extrapolation (red line) is employed to determine the current phase, and the phase offset (Equation S10-1) is added to the current phase to predict the future phase.

Autoregression with Forecast Padding (ARFP)

In the conventional AR method, the edge portion of the filtered signal near $t =$

$0$ is removed due to the filter edge effect. However, this removal may lead to information loss around the edge as well as an extended prediction period. To preserve the information contained in the edge portion, ARFP relies on the raw EEG up to the current sample ($t \leq 0$) and only uses the forecast signal for the future part ($t > 0$). The overall process is illustrated in Figure S10e, which is slightly modified from the concept of our previous work [21]. First, the signal is zero-phase bandpass filtered within 8-13 Hz; then, the edges at both ends (shaded gray regions) are removed due to the filter edge effect. The remaining part (blue line) is used to fit the autoregressive (AR) model. The fitted AR coefficients are then used to iteratively predict future samples by linearly combining the previous samples. The forecast signal at $t < 0$ (red dashed line) is discarded, and the future portion (red solid line) is concatenated with the raw EEG signal (black line). For smooth concatenation, the two signals are shifted to the same baseline level around $t = 0$ before concatenation. The concatenated signal is zero-phase bandpass filtered again in the alpha band, and the Hilbert transform is applied to estimate the current phase at $t = 0$. Finally, the phase offset (Equation S10-1) is added to the estimated current phase to predict future phases.

ETP, FFT, and PR with Forecast Padding (ETPFP, FFTFP, and PRFP)

The predicted phases of the original algorithms are converted to the forecast signal, namely the padding, as follows:

$$\text{padding}(t) = A\cos\phi_p(t) \quad \text{(S10-2)}$$

. Here, $\phi_p(t)$ is the predicted phase and $A$ is the mean instantaneous amplitude of the filtered signal in the first convolution, with the first and last 330 points excluded due to filter edge effect. The padded signal (black and red) is filtered again to extract the current phase at $t = 0$ and phase offset (Equation S10-1) is added to predict future phases.

### S.11 Online Experiments

Human Participants

The study was approved by Institutional Review Board of Cheng Hsin General Hospital. All participants received informed consent prior to participation. Twenty subjects (8 males and 12 females; mean age ± SD: 23.6 ± 4.7 years; age range: 20-40 years) were recruited according to the following inclusion criteria.

1. Healthy, right-handed subjects with normal motor function
2. No history of neurological or psychiatric disorders or related drug use
3. Age range from 18 to 70 years old

#### System Configuration

32-channel EEG (10-20 system, Fz as reference electrode) was collected by an amplifier (actiCHamp, Brain Products GmbH, Germany) with 1kHz sampling rate and was sent to a host PC (Microsoft Windows 11, 4 cores, 2.42 GHz CPU, 16GB RAM) running the signal monitoring software (BrainVision Recorder 1.25.0101, Brain Products GmbH, Germany). The EEG was then streamed to a processing computer (Microsoft Windows 11, 6 cores, 2.59 GHz CPU, 8 GB RAM) running the phase prediction algorithms (written in MATLAB R2022b) via remote data access (RDA). In the real-pulse session, RS232 serial communication was established between the processing computer and the TMS machine (Rapid$^2$, Magstim Co., Ltd., UK) using the MAGIC toolbox [22]. The processing computer issues a trigger command once the predefined target phase is met. The prediction of future phases takes place every time the new data chunk (20 samples) came in and was suspended in the 5-second periods after stimulation due to the large artifact. In the fake-pulse session, the TMS trigger cable was unplugged, and the event marker generator (TriggerBox Plus, Brain Products GmbH, Germany) was connected between the processing computer and the actiCHamp amplifier. Markers were sent to the amplifier and stored with the EEG data.

In order to control the experiment time, if the algorithm could not detect the target phase in 10 seconds, corresponding to 500 prediction attempts, the TMS trigger or the marker was sent forcefully and the system moved on to the next stimulus even if the predicted phase was not in the trigger range. These forcefully terminated trials are hereafter referred to as "skipped trials".

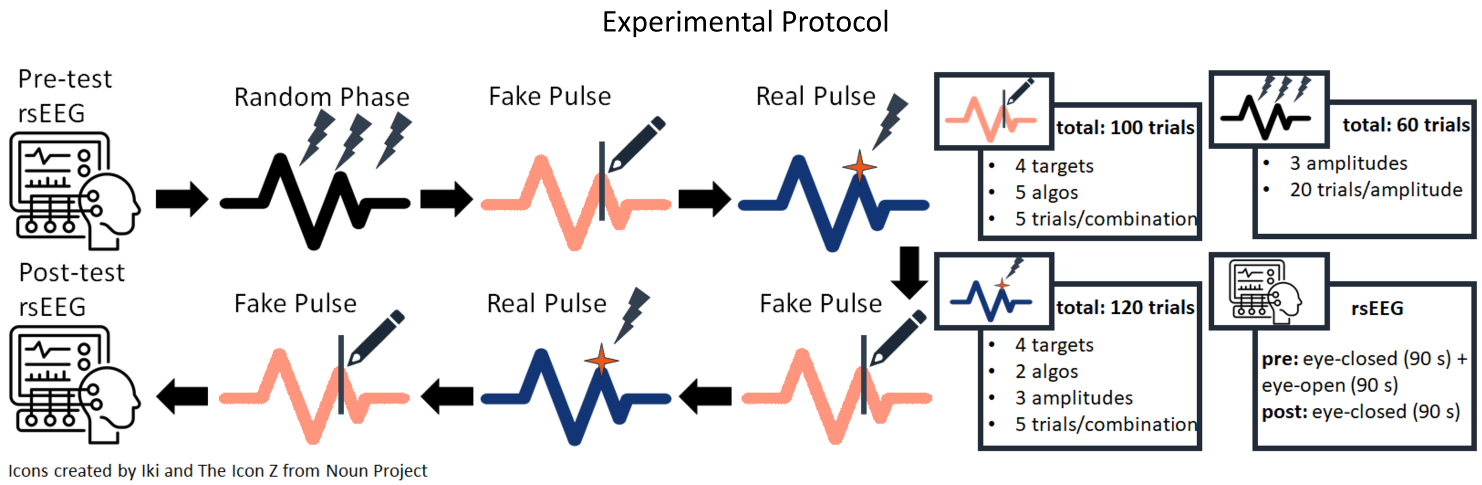


Figure S11: Resting-state EEG (rsEEG) was recorded at the beginning and the end of the experiments. The eye-open rsEEG at the beginnig was used by genetic algorithm to optimize the parameters of the phase prediction algorithms. In the random TMS session, the inter-pulse interval was uniformly distributed in 4.95 to 5.05 seconds. The stimulation amplitude was set to be 1, 1.3, and 1.6 times of RMT with 5 pulses delivered for each amplitude, summing up to 60 stimuli. The trials with different amplitudes were randomly interleaved. In fake-pulse session, 4 target phases (0, 90, 180, and 270 degrees) and 5 algorithms were randomly interleaved. Each target-algorithm combination was associated with 5 stimuli, in which the event markers were sent to the EEG amplifier if the prediction at $t = \mathbb{E}(\tau_{all})$ was in the trigger range (target $\pm$ 0.2 radians). In real-pulse session, 4 target phases (0, 90, 180, and 270 degrees), 2 algorithms (AR and ARFP) and 3 stimulation amplitudes (1, 1.3, and 1.6 times of RMT) were randomly interleaved. Each target-algorithm-amplitude combination was associated with 5 stimuli, in which actual TMS pulse was triggered if the prediction at $t = \mathbb{E}(\tau_{all})$ was in the trigger range (target $\pm$ 0.2 radians). Here, $\mathbb{E}$ denotes the expected value approximated by sample mean.

The experiment procedure is illustrated in Figure S11. Each subject underwent 2 resting-state EEG (rsEEG) sessions at the beginning and end of the experiment, 1 random TMS sessions, 3 fake-pulse sessions, and 2 real-pulse sessions.

##### rsEEG Session and RMT Measurement

In the first rsEEG session, the subject was asked to remain still, open the eyes for 90 seconds and then close the eyes for another 90 seconds. Eye-open rsEEG was used to optimize the parameters (Supplementary S.2) and phase offset (Equation S10-1) of the phase prediction algorithms. Immediately after the first rsEEG session, the subject's resting motor threshold (RMT) was estimated. During RMT measurement, electrodes were attached to the abductor pollicis brevis (APB) and the first dorsal interosseous (FDI) muscles in a belly-tendon montage, and the TMS coil was positioned over the motor hotspot (the site that generates the highest motor-evoked potential) [23]. The EMG signals were pre-amplified by a differential amplifier (BIP2AUX adapter, Brain Products GmbH, Germany) and then transmitted to the auxiliary channel of the actiCHamp amplifier. RMT was estimated using 30 pulses with the threshold hunting approach [24, 25]. The lower RMT value of the APB and FDI muscles was selected and capped at 62.5% of the maximum stimulator output (MSO) if it exceeded this threshold. Stimulation amplitudes were set to 1, 1.3, and 1.6 times this value. The second rsEEG session was arranged at the end of the experiment, where subjects were asked to remain still and close their eyes, and 90 seconds of rsEEG were recorded.

##### Random TMS Session

In random TMS session, TMS pulses were delivered without phase synchronization. The three amplitudes randomly took place and each was associated with five stimuli.

##### Fake-Pulse Session

In fake-pulse session, 5 algorithms (AR, ETP, FFT, PR, and ARFP) and 4 target phases (0°, 90°, 180°, and 270°) were tested and event markers were delivered if the predicted phase at $\mathrm{t} = \mathbb{E}(\tau_{all})$ was in the trigger range ($\tau_{all}$ varied between trials, so the sample mean $\mathbb{E}(\tau_{all})$ was used). Note that the prediction length was set to $\mathbb{E}(\tau_{all})$ rather than $\mathbb{E}(\tau_{all}^{fake})$ because we were interested in the accuracy at the actual pulse location.

##### Real-Pulse Session

In real-pulse session, 2 algorithms (AR and ARFP) and 4 target phases (0°, 90°, 180°, and 270°) were included and TMS was triggered if the predicted phase at $\mathrm{t} = \mathbb{E}(\tau_{all})$ was in the trigger range.

### S.12 Data Preprocessing

Online Experiments

The multichannel EEG was converted to a single-channel C3-laplacian montage [2, 26] for use in both the online experiments and subsequent offline analysis. During the algorithm calibration stage, the ground truth was obtained by applying zero-phase filtering and the Hilbert transform to the continuous laplacian signal. The EEG and corresponding ground truth were divided into 26 overlapping epochs (4-second epochs, stride = 1 second) for parameter optimization (see Supplementary S.2). During the online experiments, the latest 2 seconds of the laplacian signal were sent to the five algorithms described in Supplementary S.10 to predict future phases. Trigger information and EEG data were stored for subsequent accuracy and robustness evaluation.

Offline Analysis

The Laplacian signals from both the fake-pulse and real-pulse sessions were used for accuracy and robustness evaluation, with exclusion criteria as described in Supplementary S.4. The filters and algorithm parameters were identical to those used in the online experiments.
